# A Distinctive Pattern of Hippocampal-Prefrontal Cortical Network Activity during Stress Predicts Learned Resistance

**DOI:** 10.1101/801365

**Authors:** Danilo Benette Marques, Rafael Naime Ruggiero, Lezio Soares Bueno-Junior, Matheus Teixeira Rossignoli, João Pereira Leite

## Abstract

The perception of control over a stressful experience may determine its impacts and generate resistance against future stressors. Although the medial prefrontal cortex (mPFC) and the hippocampus are implicated in the encoding of stressor controllability, the neural dynamics underlying this process are unknown. Here, we recorded CA1 and mPFC neural activities in rats during the exposure to controllable, uncontrollable, or no shocks, and investigated electrophysiological predictors of escape performance upon exposure to subsequent uncontrollable shocks. We were able to accurately discriminate stressed from non-stressed animals and predict resistant or helpless individuals based on neural oscillatory dynamics. We identified a pattern of enhanced CA1-mPFC theta power, synchrony, cross-frequency interaction, and neuronal coupling that strongly predicted learned resistance, and that was lacking in helpless individuals. Our findings suggest that hippocampal-prefrontal network theta activity supports cognitive mechanisms of stress coping, and its impairment may underlie vulnerability to stress-related disorders.

## Introduction

The perception of control over a stressful experience is critical to determine its impacts on the individual (Southwick and Charney, 2012). Learning that adverse situations are controllable or uncontrollable drives the acquisition of long-term resistance or vulnerability against future stressors, respectively (Maier and Seligman, 1976, 2016). Rats exposed to uncontrollable inescapable shocks (IS) exhibit deficient escape learning, potentiated anxiety, delayed fear extinction, disrupted serotonergic transmission, and impaired neural plasticity (Amat et al., 1998, 2005, 2006, 2010; Baratta et al., 2007; Maier and Watkins, 1998, 2005; Seligman and Beagley, 1975; Seligman and Maier, 1967; Shors et al., 1989, 2007). The failure to escape after uncontrollable aversive events was classically named “learned helplessness” (Seligman and Maier, 1967), and numerous biological effects parallel to this behavior bear translational validity with clinical depression and anxiety disorders (Maier and Watkins, 1998; Pryce et al., 2011; Seligman, 1975; Vollmayr and Gass, 2013; Willner, 1984). In contrast, subjects exposed to equivalent controllable escapable shocks (ES) do not present such alterations, and in addition become resistant against subsequent IS (Amat et al., 2006; Baratta et al., 2007; Maier, 2015). Therefore, investigating the neurobiology of stressor controllability may elucidate mechanisms of resistance to stress-related disorders and their treatments.

Foundational work from Maier *et al.* demonstrated that inhibition of mPFC activity and plasticity abolishes the acute and long-term protective effects of stressor controllability in a wide range of behavioral and physiological outcomes (Amat et al., 2005, 2006; Baratta et al., 2008; Maier, 2015; Maier and Watkins, 2010; Maier et al., 2006). Many reports also show that the hippocampus is differentially sensitive to controllable and uncontrollable stress (Amat et al., 1998; Balleine and Curthoys, 1991; Hadad-Ophir et al., 2017; Shors et al., 1989, 2007), and that intra-hippocampal administration of antidepressants prevents the development of helplessness after IS (Joca et al., 2003). Moreover, hippocampal-prefrontal cortical functional connectivity (Gordon, 2011; Thierry et al., 2000) is modulated by both stress and antidepressants, and participates in both emotional and higher-order cognitive processes that are dysfunctional in stress-related disorders (Godsil et al., 2013; Jay et al., 2004; Rocher et al., 2004; Zheng and Zhang, 2015). Although it has been well established that mPFC and hippocampus play critical roles in encoding stressor controllability, the neural dynamics underlying this process remain unknown.

Here we hypothesized that the encoding of stressor controllability and uncontrollability would be associated with distinct patterns of neural activity related to CA1 and mPFC interaction during stress. To test this hypothesis, we simultaneously recorded CA1 and mPFC local field potentials (LFP) and single-unit activity in rats during the exposure to either ES, yoked IS, or no shocks (NS), all of which signaled by identical conditioned stimuli (CS). Then, we explored electrophysiological predictors of learned resistance or helplessness to subsequent uncontrollable shocks, as determined by later escape performance. We found an unprecedented association between stressor controllability and enhanced CA1-mPFC theta power and synchrony, as well as mPFC local theta phase coupling to both fast oscillations and neuronal firing during the anticipation of aversive stimuli. We were also able to implement a computational model that accurately discriminated stressed from non-stressed animals and predicted resistant individuals based solely on oscillatory dynamics. Our results indicate that mPFC theta activity entrained by CA1 underlies the encoding of stressor controllability, and the lack of this protective activity may allow the development of helplessness in the face of severe stress.

## Results

### Differential Engagement of Theta Oscillations during the Expectation of Controllable and Uncontrollable Stress

To identify neural correlates of stressor controllability and uncontrollability within CA1 and mPFC neural dynamics, we submitted rats to controllable ES, yoked uncontrollable IS or NS - all of which signaled by conditioned stimuli (CS) - in a custom shuttle box that allowed electrophysiological recording and footshock delivery (Figure 1A, see Methods). ES rats (N = 11) could escape footshocks by jumping a short wall between compartments, while yoked IS counterparts (N = 9) received equivalent uncontrollable shocks. All shocks were preceded by a conditioned light stimulus (CS^+^), that was presented alone (CS^-^) for NS rats (N = 7). We hypothesized that the neural activity during the anticipation of aversive stimuli (CS period) would depend on the expectation of controllability or uncontrollability. Initially, we confirmed that all ES animals learned the escape response on day 1 (Figure 1B). To evaluate if this initial exposure would generate resistance to future stressors, we exposed both ES and IS animals to uncontrollable shocks of fixed duration (10 s) in the same apparatus on day 2. Then on day 3, i.e., the test session, we measured the escape performance of all animals (Figure 1A). Our results show that both the mean latency to escape and the number of failures were equivalent between ES and NS, but lower in ES than IS (latency: one-way ANOVA F(2,24) = 4.77, p = 0.01; post hoc Fisher’s LSD test: ES vs. NS: t(24) = 0.95, p = 0.36; ES vs. IS: t(24) = 2.28, p = 0.03; failures: Kruskal-Wallis H(3) = 7.41, p = 0.02; post hoc Wilcoxon rank-sum test: ES vs. NS: U = 33, p = 0.62; ES vs. IS: U = 22, p = 0.03; Figure 1C). This observation confirms the role of controllability in mediating resistance. We then categorized resistant (R) versus helpless (H) individuals through cluster analysis based on the similarity with NS behavior (Figure 1D). Previous exposure to ES generated a greater proportion of R individuals (72%, N = 8/11) not significantly different from NS (X^2^(1, N = 18) = 2.29, p = 0.13), while IS generated a greater proportion of H individuals (66%, N = 6/9) significantly different from NS (X^2^(1, N = 16) = 7.46, p = 0.006) (Figure 1E). Notoriously, resistant and helpless animals showed clearly distinct behavioral profiles in the test session (latency: F(4,22) = 18, p < 0.0001; number of failures: H(5) = 18.57, p = 0.001) (Figure 1F-G). Also noteworthy, we identified smaller subsamples of H individuals that underwent ES (H-ES, N = 3) and R individuals that underwent IS (R-IS, N = 3) (Figure 1A, E). We reasoned, though, that the neural correlates of controllability and uncontrollability would be best represented in ES animals that became R (R-ES, N = 8) and IS animals that became H (H-IS, N = 6) respectively. Then, to further validate these neural signatures (see Discussion), we examined if they could also predict behavior in H-ES and R-IS animals, or correlate with subsequent escape performance considering the entire sample of stressed animals.

**Figure 1.**
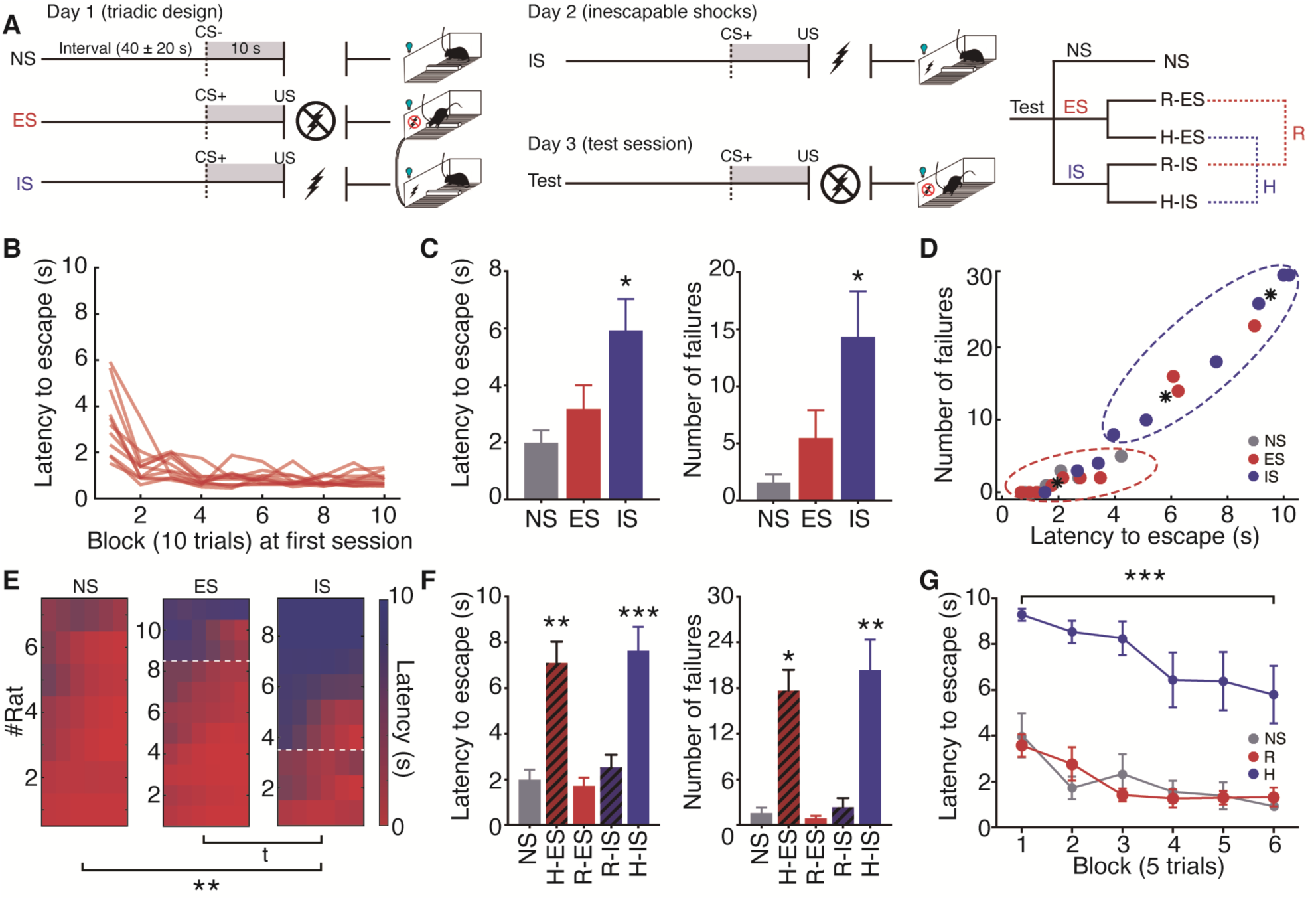
The Triadic Design of Stress Exposure Induces Learned Resistance and Helplessness. (A) Triadic design of stress exposure (day 1) and “immunization” protocol. Previously stressed rats were submitted to IS (day 2). Escape performance was evaluated for all animals in the test session (day 3). See Methods. (B) All ES animals learned the escape response at the first exposure (day 1). (C) IS animals presented greater mean latency to escape (*p < 0.05, Fisher’s LSD test) and number of failures (*p < 0.05, Wilcoxon rank-sum test) in the test session. (D) K-means clustering of resistant and helpless individuals. (E) Latency to escape across blocks (of 5 trials) (x-axes) in the test session for each rat (y-axes). Helpless animals are above the dashed line. IS induced higher proportion of helpless animals (*p < 0.05, Chi-squared test). (F) Resistant and helpless animals exhibited markedly distinct behaviors in the test session (latency to escape: *p < 0.01, Fisher’s LSD test; number of failures: *p < 0.05, Wilcoxon rank-rum test). (G) R and NS animals showed identical behavior in the test session. H animals exhibited delayed escape learning. Here and on: ES = escapable shocks, IS = inescapable shocks, NS = no shocks, R = resistant, H = helpless, CS = conditioned stimulus (light), US = unconditioned stimulus (shock). Error bars represent the mean ± SEM.

We found significant effects of stressor controllability in CS^+^ event-related power perturbation (ERPP) that were exclusive to the theta band (5-10 Hz) in both mPFC (F(2,18) = 11.25, p = 0.0007; post hoc: R-ES vs. H-IS: t(18) = 4.51, p = 0.0003) and CA1 (F(2,17) = 4.89, p = 0.02; post hoc: R-ES vs. H-IS: t(17) = 3.11, p = 0.006) (Figure 2). Moreover, the CS^+^ modulations of theta power were the opposite between R-ES and H-IS groups: while R-ES animals presented a mean increase of theta power in both mPFC (0.85 ± 0.19 dB) and CA1 (0.60 ± 0.39 dB), H-IS animals exhibited a decrease (mPFC: - 0.47 ± 0.26 dB; CA1: −0.75 ± 0.19 dB) (Figure 2C-D). Furthermore, we found a negative correlation between theta ERPP on day 1 and latency to escape in the test session (day 3) across all stressed animals in both brain regions. This correlation was much stronger in the mPFC (r(18) = −0.77 p < 0.0001) than CA1 (r(18) = −0.58, p = 0.006) (Figure 2E). Remarkably, R-IS and H-ES subsamples exhibited a power perturbation profile more closely linked to their behavioral outcomes (R vs. H) than the programmed exposure (ES vs. IS) they were going through (Figure S2). In addition, CS^+^ profoundly decreased power in a wide range of frequencies from delta to alpha/beta bands (1-30 Hz), regardless of the degree of control (See also Figure S3). Then, we found that the association between stimulus-triggered CA1 theta power and stress control selectively occurred during CS^+^, while the periods following the interruption of the unconditioned stimuli (US, footshocks, see Methods) were marked by brief increases in theta peak frequency, not power, irrespective of their controllability (Figure S4). Taking these differences into account, we show through principal component analysis (PCA) that CA1 theta power-frequency modulation across stimuli can map stress-related behavioral and cognitive states of anticipation (CS^+^) reaction (US), and controllability (CS^+^ in R-ES vs. H-IS) (Figure S4F-H). Although we found a suppression of mPFC theta power during CS^+^ associated with helplessness, H-IS exhibited stronger theta phase resetting to CS^+^ onset (Figure S5). Overall, these results demonstrate that the sustained engagement of CA1 and mPFC theta power during the anticipation of aversive stimuli depends on the expectation of control, and that this activity is related to the acquisition of resistance to subsequent stressors.

**Figure 2.**
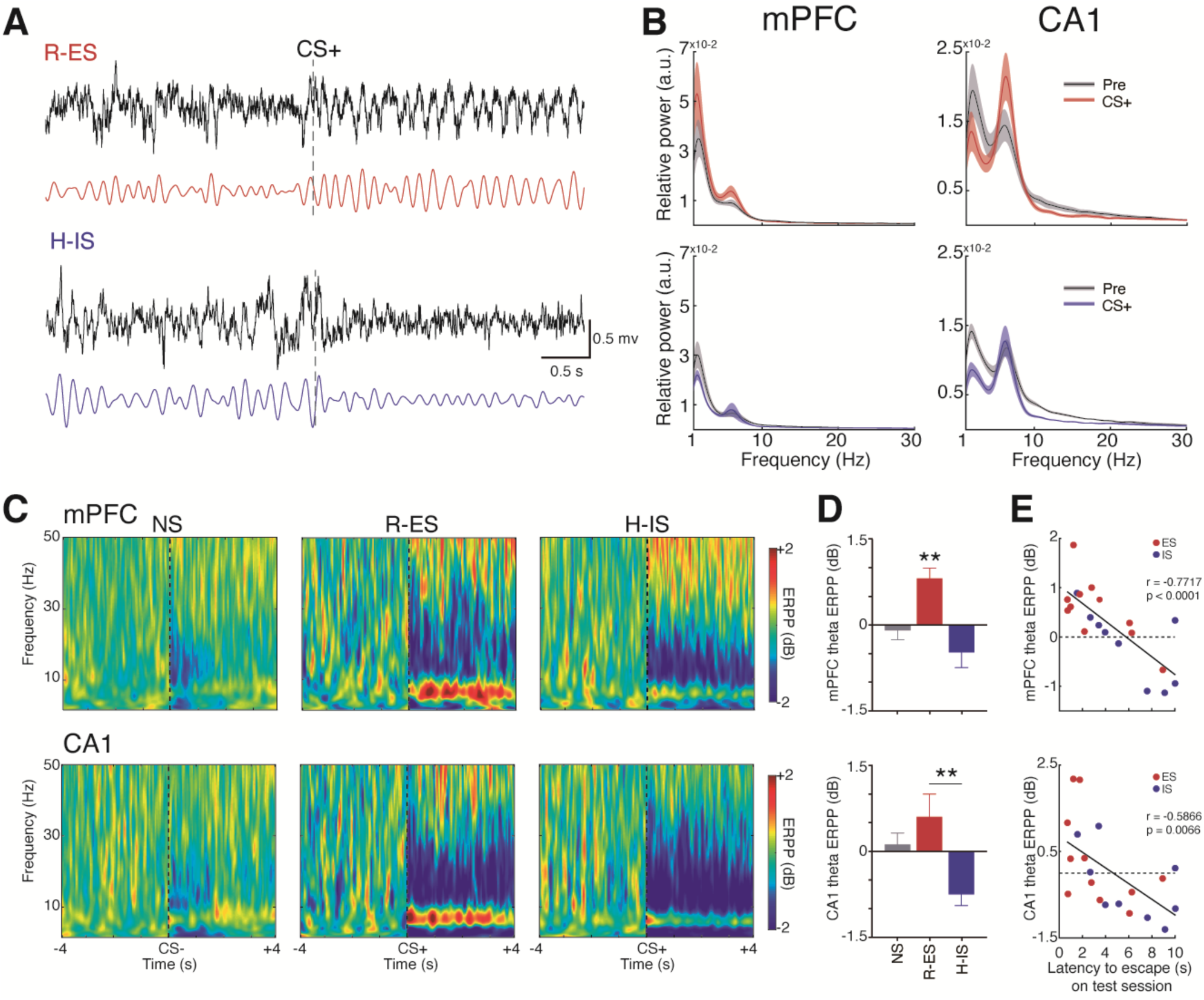
Differential Engagement of Theta Oscillations during the Expectation of Controllable and Uncontrollable Stress. (A) Representative traces of CA1 LFPs (black) and theta-filtered signals (colored) responses to CS^+^ preceding controllable (top) and uncontrollable (bottom) shocks. Note the opposing effects on amplitude. (B) mPFC and CA1 relative power spectral densities (mean ± SEM) showing an increased theta (5-10 Hz) power in R-ES during CS^+^ that is absent in H-IS. (C) Average spectrograms showing a unique distinction between R-ES and H-IS responses in the theta band. Note that CS^+^ promotes a wide-spectrum power reduction (1-30 Hz) regardless of controllability. (D) Opposed CS^+^-related modulations of theta power in R-ES (increase) and H-IS (decrease). (F) mPFC and CA1 theta power modulations in the first session (day 1) correlate with later escape performance (day 3). All animals under stress are included. ERPP = event-related power perturbation. *p < 0.05, Fisher’s LSD test. Here and on: shaded lines represent the mean ± SEM. See also Figures S2-5.

### A Distinctive Pattern of mPFC Spectral Perturbation during Stress Correlates with Later Escape Learning

After describing the relationship between mPFC theta and the expectation of control, we decided to investigate how power perturbations in a broader spectrum could be related with escape learning during new aversive events. As previously shown in Figure 1, we noticed more variability in escape latency in the final trials of the test session (Figure 1G), consistently with the observation that some H animals showed either delayed learning or no learning at all (Figure 1E). Since impaired escape learning is an essential feature of helplessness, we evaluated whether ERPP frequencies correlate with escape performance across trials of the test session. Initially, we confirmed that theta-band ERPP in the mPFC had the strongest correlation with the mean escape performance (Figure 3A). Then, we found that this correlation was strong throughout the entire test session, whereas in CA1 it was confined to the initial trials (Figure 3B-C). This finding supports a link between mPFC activity – and perhaps mPFC-dependent cognition/plasticity – and the preserved ability to learn to escape.

**Figure 3.**
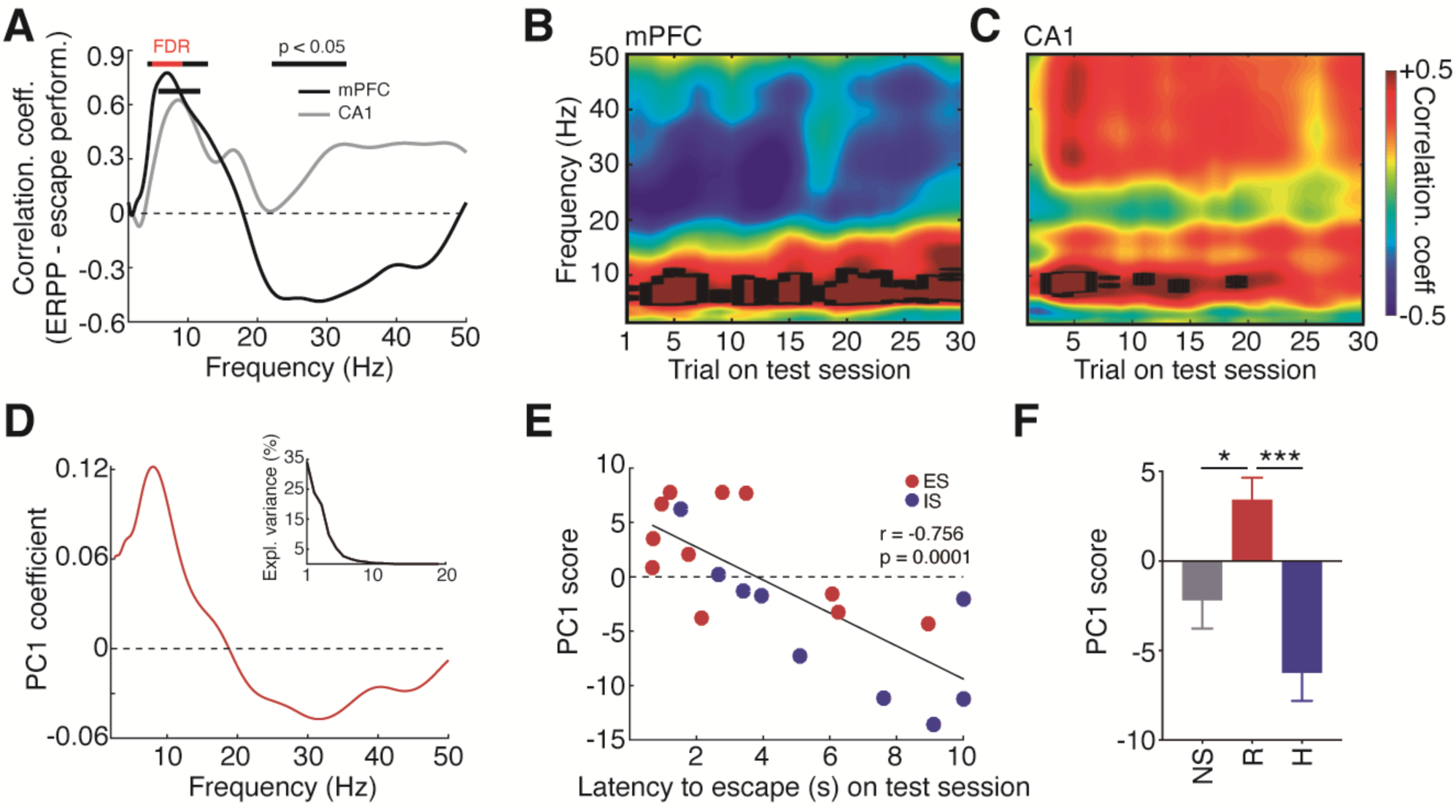
A Distinctive Pattern of mPFC Spectral Perturbation during Stress Correlates with Later Escape Learning. (A) Spectrum of power modulation correlations (Pearson’s r coefficient) to subsequent escape performance revealed specific patterns. Theta band presented the greatest correlation, which was stronger in the mPFC than CA1. Escape performance was distinctively associated with low gamma band (30-50 Hz) in CA1 (positive) and mPFC (negative). Black line: p < 0.05, red line = p < false discovery rate corrected, Pearson’s correlations. (B and C) mPFC, but not CA1, presented a strong correlation with escape performance in the final trials of the test session, indicating a preserved ability of escape learning. Black lines = p < false discovery rate corrected. (D) mPFC power perturbation PC1 coefficients revealed the same pattern as the mPFC spectrum of correlations. (E) PC1 scores showed greater correlation with escape than each frequency separately. (F) Significant modulation of PC1 scores in R animals. *p < 0.05, Fisher’s LSD test.

Despite the stronger correlations in the theta band, other frequencies also showed correlations with behavior (Figure 3A-B). We noticed that some frequencies, such as in the low gamma range (30-50 Hz), exhibited distinct associations with escape learning depending on the brain region – positive in CA1 and negative in mPFC (see also Figure S3). These observations led us to hypothesize the presence of a wide-spectrum collective pattern of cross-frequency power relationships that could be linked to the behavioral outcomes of stress. We explored this using PCA of the power-perturbed spectra (2.5-50 Hz, heuristically-determined range) from animals under stress (N = 20). Surprisingly, the PC1 coefficients (Figure 3D) – which collectively account for the maximum data variance (35% explained variance; see inset of Figure 3D) - showed basically the same pattern as the spectrum of correlations with escape performance (Figure 3A). Additionally, PC1 scores showed stronger correlation with subsequent escape latency (r(18) = −0.75, p = 0.0001; Figure 3E) than each frequency separately (see Figure 3A), and were significantly greater in R animals (F(2,24) = 12.57, p = 0.0002; R vs. H t(24) = 4.97, p < 0.0001; R vs. NS (t(24) = 2.68, p = 0.01; H vs. NS t(24) = 1.85, p = 0.07; Figure 3F). In summary, these results indicate that variations across a wide spectrum of stress-related mPFC power perturbation are associated with helplessness or resistance.

### Enhanced CA1 to mPFC Theta Synchrony during Stress Correlates with Learned Resistance

To assess more directly the functional connectivity between CA1 and mPFC, we calculated their phase coherence. The mean phase coherence (MPC) spectra revealed a unique increase within the theta band during CS^+^ in both R-ES and H-IS, but not NS animals (Figure 4A). This effect was observed to be consistent across all R-ES and H-IS subjects (see insets of Figure 4A). We also show that although theta mean resultant length (MRL) did not differ between groups during the pre-CS period, it was significantly modulated by both CS^+^ and the expected degree of control (two-way ANOVA stressor controllability x period interaction F(2,34) = 7.73, p = 0.001) (Figure 4B). Still in this analysis, we found that theta MRL in both R-ES and H-IS were greater than NS (t(34) = 6.01, p < 0.0001; t(34) = 2.82, p = 0.007), and this effect was even greater in R-ES than H-IS (t(34) = 2.99, p = 0.005) (Figure 4B). The directionality of theta LFP between regions was examined through cross-correlation analysis, and we showed that CA1 theta signals constantly led the mPFC by ∼49 ms during stress (peak of the averaged cross-correlation = −49 ms, Figure 4C; bin with the maximum number of lags = −48.14 ms; Figure 4D). To further illustrate these findings, we plotted polar histograms of theta phase differences from representative animals of each behavioral category (Figure 4E). In NS, the angles were homogeneously distributed. In stressed animals, on the other hand, these distributions were centralized around a certain angle (∼130°, consistent with the cross-correlation lags), which became even more concentrated during CS^+^, especially in the representative R-ES individual (Figure 4E). Moreover, we estimated time-frequency CS^+^-related modulation of phase coherence, and we found a stronger CA1-mPFC synchronization in R-ES than H-IS (F(2,17) = 11.14, p = 0.0080; t(15) = 3.21, p = 0.005; Figure 4F-G). Finally, CA1-mPFC theta MPC during CS^+^ was strongly correlated with the mean escape latency in the test session (r(18) = −0.70, p = 0.0005; Figure 4H). In addition to corroborating previous reports that CA1 theta synchronizes with the mPFC during the anticipation of aversive stimuli (Seidenbecher et al., 2003), these results demonstrate that the strength of this synchronization is associated with controllability and learned resistance.

**Figure 4.**
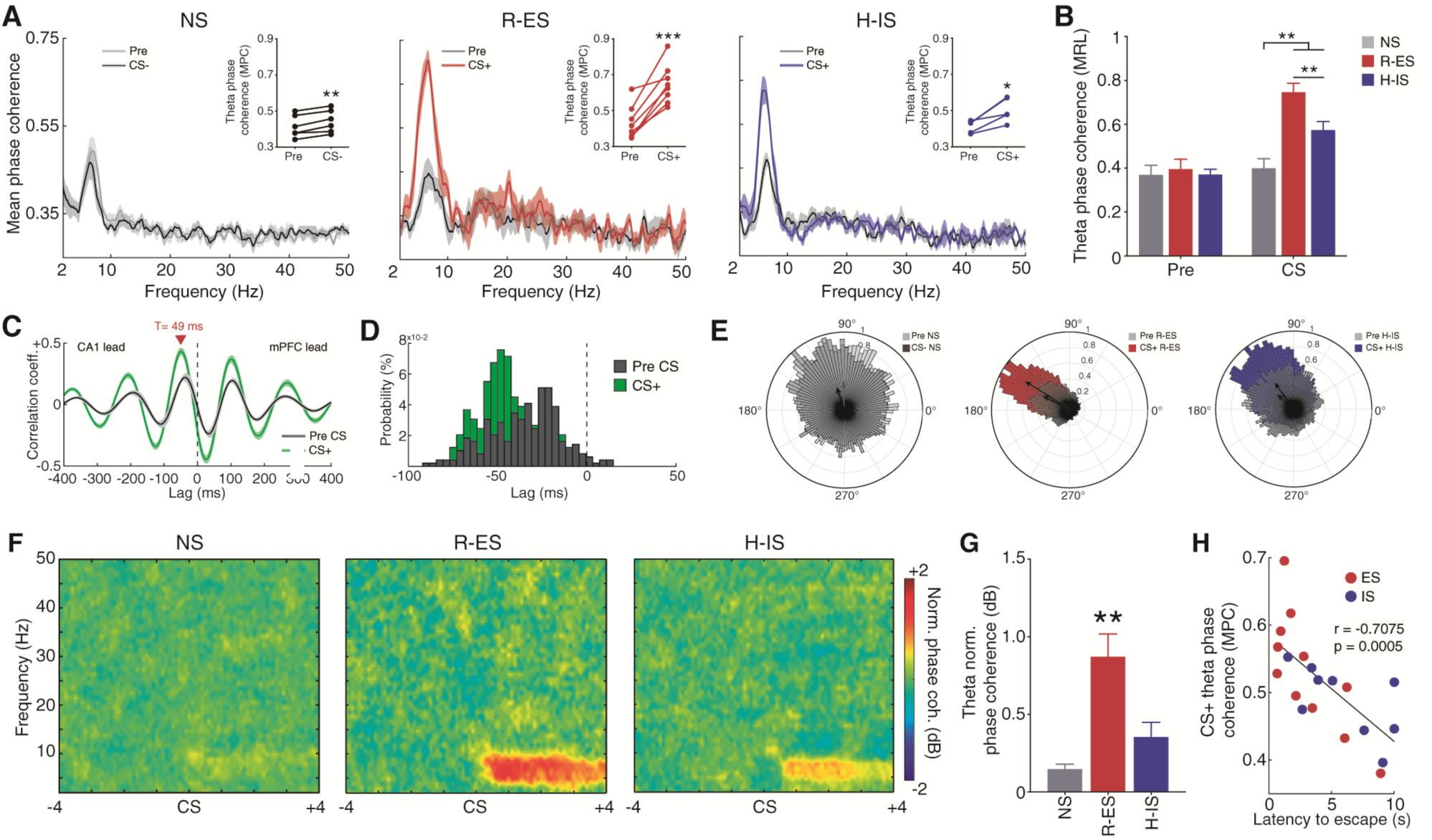
Enhanced CA1 to mPFC Theta Synchrony during Stress Correlates with Learned Resistance. (A) Mean phase coherence spectra showing a specific modulation of theta band by CS^+^. Insets show that all animals under stress presented theta synchronization (*p < 0.05, paired t-test or signed rank test). (B) Greater theta mean resultant length during CS^+^, particularly stronger in R-ES. There were no differences in the pre-CS period. (C to D) CA1 theta consistently preceded the mPFC in ∼49 ms. (C) Average cross-correlation coefficients for all trials (N = 489) of animals under stress and (D) distribution of lags of coefficient peaks. (E) Representative polar histogram of CA1 to mPFC theta phase differences showing homogenous distribution in the NS animal, consistent phase concentration in both stressed animals around 130° (∼49 ms), and even greater concentration in the R-ES animal. Vector sizes indicate the mean resultant length. (F) Average spectrograms of event-related phase coherence perturbation. (G) R-ES presented significantly greater phase coherence perturbation. (H) Theta mean phase coherence during CS^+^ presented a strong correlation with later escape performance.

### Prestimulus Theta-Gamma Phase-Amplitude Coupling in the mPFC is Associated with Stress Resistance and CS^+^-Evoked Theta Power Modulation

The findings reported so far show that the network activity during CS^+^ is highly associated with the expected degree of control over upcoming aversive stimuli. However, especially in the pre-CS period we observed a distinctively strong theta-gamma phase-amplitude coupling (Figure 5A) in resistant individuals. From the comodulation maps shown in Figure 5B, we observed that phase-amplitude coupling was specific to theta phases (4-10 Hz) and high gamma (80-110 Hz) amplitudes (Figure 5B-C). Curiously, mPFC pre-CS theta-high gamma modulation index (MI) was more effective in discriminating R from H individuals (H(2) = 9.14, p = 0.01; R vs. H: U = 20, p = 0.02; AUC = 0.79; Figure 5D) than during the CS period (AUC = 0.64). Conversely, CA1 showed no evidence of associations between theta-high gamma coupling with either stress or behavior (CA1 pre-CS MI one-way ANOVA F(2,23) = 0.58, p = 0.56; Figure 5E). Theta-gamma MI has been shown to correlate with ongoing theta power (Tort et al., 2008). Consistently, we found such a correlation, but interestingly pre-CS MI was more strongly correlated with CS^+^ theta power (r_s_(18) = 0.69, p = 0.0007) than with pre-CS theta power (r_s_(18) = 0.67, p = 0.001) (Figure 5F). In fact, pre-CS MI also correlated with theta ERPP (r_s_(18) = 0.64, p = 0.002; Figure 5G). These findings indicate that basal theta-high gamma MI in the mPFC is associated with stress resistance and controllability-related patterns of CS^+^-evoked theta modulation.

**Figure 5.**
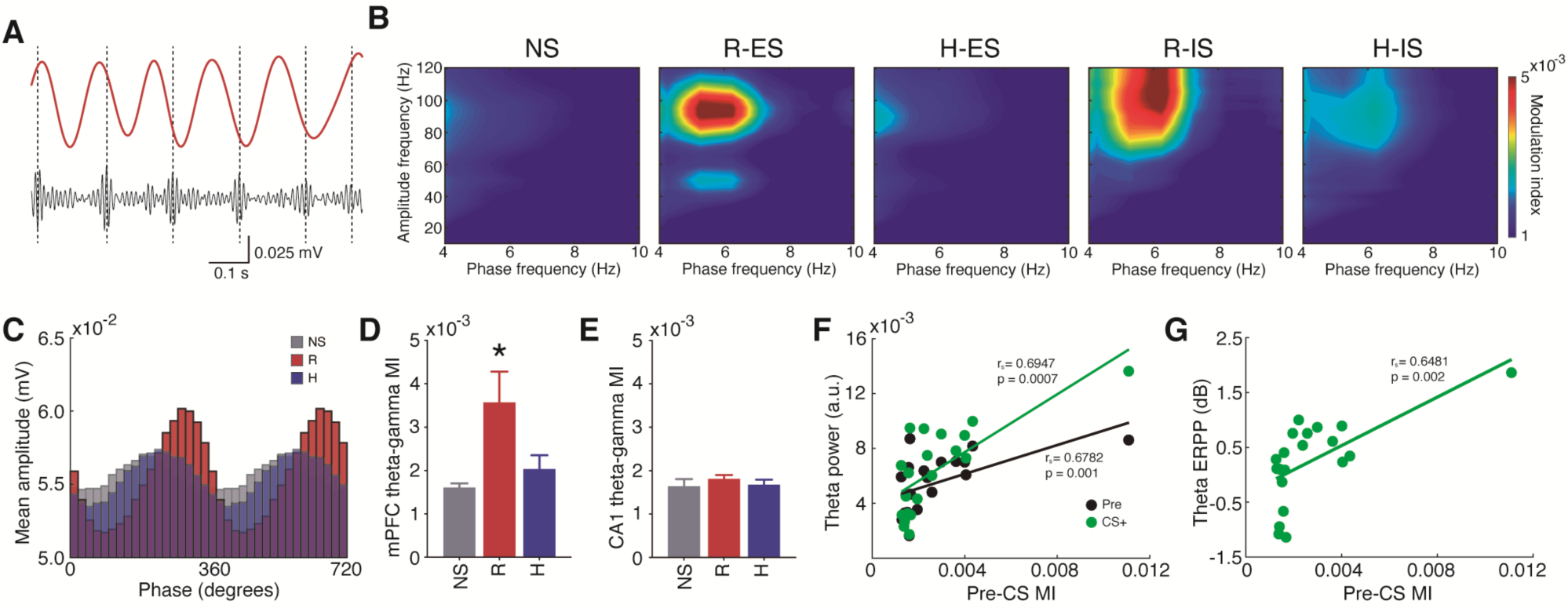
Prestimulus Theta-Gamma Phase-Amplitude Coupling in the mPFC is Associated with Stress Resistance and CS^+^-Evoked Theta Modulation. (A) Representative traces of mPFC theta (4-10 Hz, red) and high gamma (80-110 Hz, black) filtered signals showing consistent phase coupling prior to CS. (B) mPFC comodulation maps showing blobs of theta-high gamma phase-amplitude coupling in R animals from both ES and IS. (C) Theta phase and high gamma amplitude histograms showing stronger coupling in R. (D to E) The association between theta-gamma coupling and stress resistance was specific to the mPFC. (D) R presented greater MI than both NS and H (*p < 0.05, Wilcoxon rank-sum test). (F to G) mPFC pre-CS^+^ MI showed (F) a stronger correlation with CS^+^ theta power (green) than pre-CS^+^ theta power (black) and (G) also correlated with CS^+^ theta power perturbation (Spearman’s correlation).

### Differential Neuronal Firing Responses to CS^+^ and US is Associated with Controllability or Helplessness

Previous studies have shown that mPFC inhibition abolishes the protective effects of stressor controllability (Amat et al., 2005, 2006). Based on these findings, Maier *et al.* hypothesized that mPFC activity would be excited under controllable stress, and either suppressed or not active under uncontrollable stress (Maier, 2015; Maier et al., 2006). Our study addressed this question by recording mPFC neuronal activity around ES and IS and, surprisingly, we found no supporting evidence for this hypothesis (Figure 6). We analyzed the firing rates of spike-sorted neurons from NS (N = 21), R-ES (N = 33), and H-IS (N = 35) (Figure 6A). We observed a higher incidence of significantly stimulus-modulated neurons in stressed than NS animals (X^2^(1, N = 110) = 14.65, p = 0.0001), which were represented as subsamples of both excited and suppressed units upon either CS^+^ or US triggering. Furthermore, differently from the hypothesis of mPFC excitability under controllable (but not uncontrollable) stress, the proportions of excited or suppressed neurons did not differ between R-ES and H-IS, regardless of the trigger (CS^+^: X^2^(2, N = 68) = 0.87, p = 0.64); US: X^2^(2, N = 68) = 0.11, p = 0.94) (Figure 6B). Finally, we separately analyzed these subsamples of neurons, and found that excitation was actually enhanced in H-IS neurons, specifically after US termination (F(2,35) = 35.91, p < 0.0001; R-ES vs. H-IS t(35) = 4.34, p = 0.0001; Figure 6C-D).

**Figure 6.**
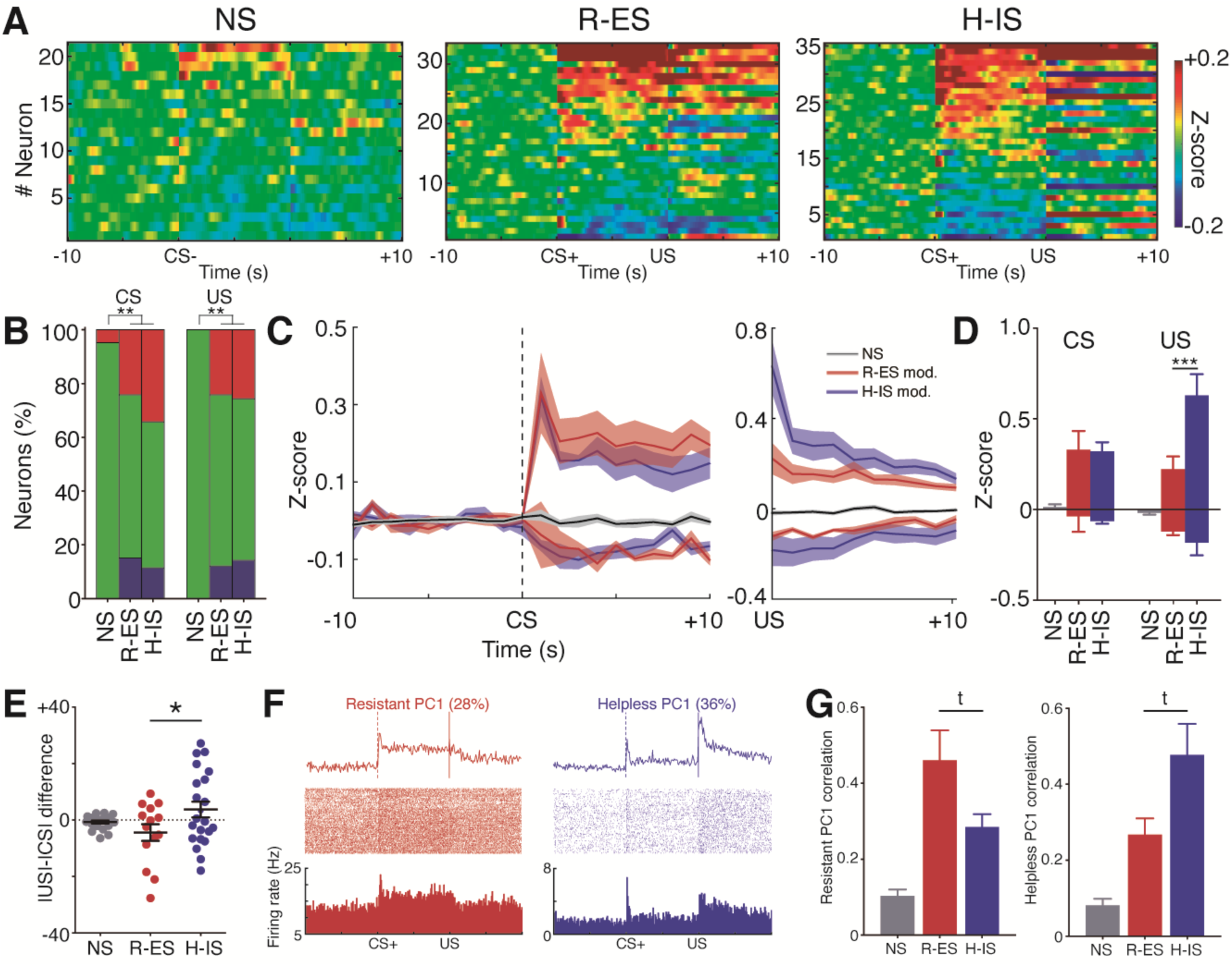
Differential Neuronal Firing Responses to CS^+^ and US is Associated with Controllability or Helplessness. (A) Firing rate modulation by CS and US (bin Z-score to the pre-CS). Neurons (y-axes) are ordered by CS modulation. There are clear distinctions between NS vs. both stressed groups, but no clear distinction between R-ES and H-IS. Additionally, note that US modulation does not correspond entirely to CS modulation. See Figure S6. (B) Both stressed groups presented more CS- and US-modulated neurons than NS but no difference was found between R-ES and H-IS (*p < 0.05, Chi-squared test). (C to D) Firing rate modulation (mean ± SEM) across neuronal categories (excited, suppressed or all NS) for each stimulus revealed that H-IS exhibited greater excitatory response after shock termination (*p < 0.05, Fisher’s LSD test). (E) Bi-directional influence of the degree of control over US-CS differences (between firing rates moduli). R-ES modulated units showed greater responses to CS^+^ than US, while H-IS showed the opposite effect (*p < 0.05, Fisher’s LSD test) (F) PC1 coefficients for all R neurons (left) and H neurons (right) with raster plots from representative neurons corresponding to each pattern. The two patterns confirm the distinction between US and CS modulation in resistance and helplessness. (G) R-ES modulated neurons tended to show greater correlation (Z-score x PC coefficients) with the *resistant PC1* (p = 0.06), while H-IS neurons tended to show greater correlation with the *helpless PC1* (p = 0.09) (^t^p < 0.1, Wilcoxon rank-sum test).

Apart from the averaged patterns of Figure 6C-D, we observed a diversity of CS^+^- and US-evoked firing modulations across individual neurons: from suppression-then-excitation to excitation-then-suppression, with a myriad of patterns in between (Figure S6). Interestingly, this diversity was much greater among stressed animals. This property suggests that the differences between CS and US may be related to the encoding of stress-relevant information at the single-neuron level (Figure S6). To approach this question, we calculated the magnitude of US-evoked versus CS-evoked responses (i.e., the difference between their absolute values) across groups (Figure 6E). This approach revealed an interesting marker of stress controllability (F(2,54) = 3.02, p = 0.056, R-ES vs. H-IS t(54) = 2.41, p = 0.019): preferential neural reactivity to CS^+^ in R-ES (−4.45 ± 2.95 Z-score) and to US in H-IS (3.70 ± 2.77 Z-score) (Figure 6E). Next, we proceeded to a deeper characterization of perievent patterns using PCA, which revealed the patterns of Figure 6F. The *resistant PC1* (28% explained variance, N = 47) was characterized by a sustained increase in firing rate during CS^+^, with a fast return to baseline after US. In contrast, the *helpless PC1* (36% explained variance, N = 42) was rather characterized by phasic responses at CS^+^ onset, but long-lived excitation at US offset. We then examined how well these perievent patterns would approximate those of CS^+^- or US-modulated neurons in R-ES versus H-IS. For this, correlations between PC1 coefficients and mean Z-scored firing rates of each neuron were converted to absolute values. We found that R-ES neurons showed greater correlation with the *resistant PC1* (H(2) = 26.28, p < 0.0001; R-ES vs. H-IS U = 97, p = 0.06), and H-IS neurons with the *helpless PC1* (H(2) = 27.04, p < 0.0001; R-ES vs. H-IS U = 102, p = 0.09) (Figure 6G). In summary, these findings demonstrate that mPFC encoding of stressor controllability and uncontrollability cannot be represented as simplistic increases or decreases in neuronal activity as previously hypothesized, but as complex perievent profiles around conditioned and unconditioned aversive stimuli.

### mPFC Neurons Modulated under Controllable Stress are Coupled with Theta Phase and Rhythm

After we found that stressor controllability was associated with mPFC theta oscillations and neuronal firing patterns, we investigated the interactions between single-neuron and oscillatory activities under controllable or uncontrollable stress. Initially, we observed that a proportion of neurons in stressed animals (22.80%, N = 13/44) showed significant phase-locked spiking to theta field potentials (Rayleigh’s Z test, p < 0.01; see representative traces in Figure 7A). In contrast, no phase-locked neurons were found in NS animals (0%, N = 0/15; X^2^(1, N = 72) = 4.17, p = 0.041; Figure 7B). Additionally, all neurons that were phase-locked to theta LFP were also modulated by stress (phase-locked modulated 100%, N = 13/13; vs. non-modulated 52%, N = 23/44; X^2^(1, N = 57) = 9.82, p = 0.001; Figure 7C). Then, we observed that modulated neurons in R-ES animals were more strongly phase-locked to mPFC theta (representative units with both phase-locking and CS^+^ responsivity are shown in Figure 7D) than those of H-IS animals (H(2) = 4.73, p = 0.093; R-ES vs. H-IS U = 45, p = 0.045; Figure 7E). We then investigated the possible association between LFP power and the spiking activity of neurons by estimating the power spectral densities (PSD) of spike-triggered averaged (STA) LFP. We found a prominent peak of STA theta power in R-ES modulated neurons (Figure 7F). In fact, STA theta power was much higher in R-ES modulated neurons than in NS neurons or H-IS modulated neurons (H(2) = 11.35, p = 0.003; NS vs. R-ES U = 24, p = 0.003; R-ES vs. H-IS U = 27, p = 0.002; Figure 7F). In addition to spike-field interactions, we also explored spike rhythmicity itself by applying PSD analysis to the binary spiking data. This approach revealed stronger theta power among stimulus-modulated neurons of R-ES than NS or H-IS (Figure 7G). Interestingly, some theta-rhythmic neurons also corresponded to the patterns of differential modulations by CS^+^ versus US we described earlier: the *resistant PC1* (greater to CS^+^); the *helpless PC1* (greater to US); and the stressor PC2 (opposed between CS^+^ and US) (Figure S7, see also Figure 6F, S6C). Altogether, our results on perievent firing rate modulation, phase-locking, and spike periodicity suggest that theta rhythm coordinates the neuronal dynamics associated with stressor controllability within the mPFC.

**Figure 7.**
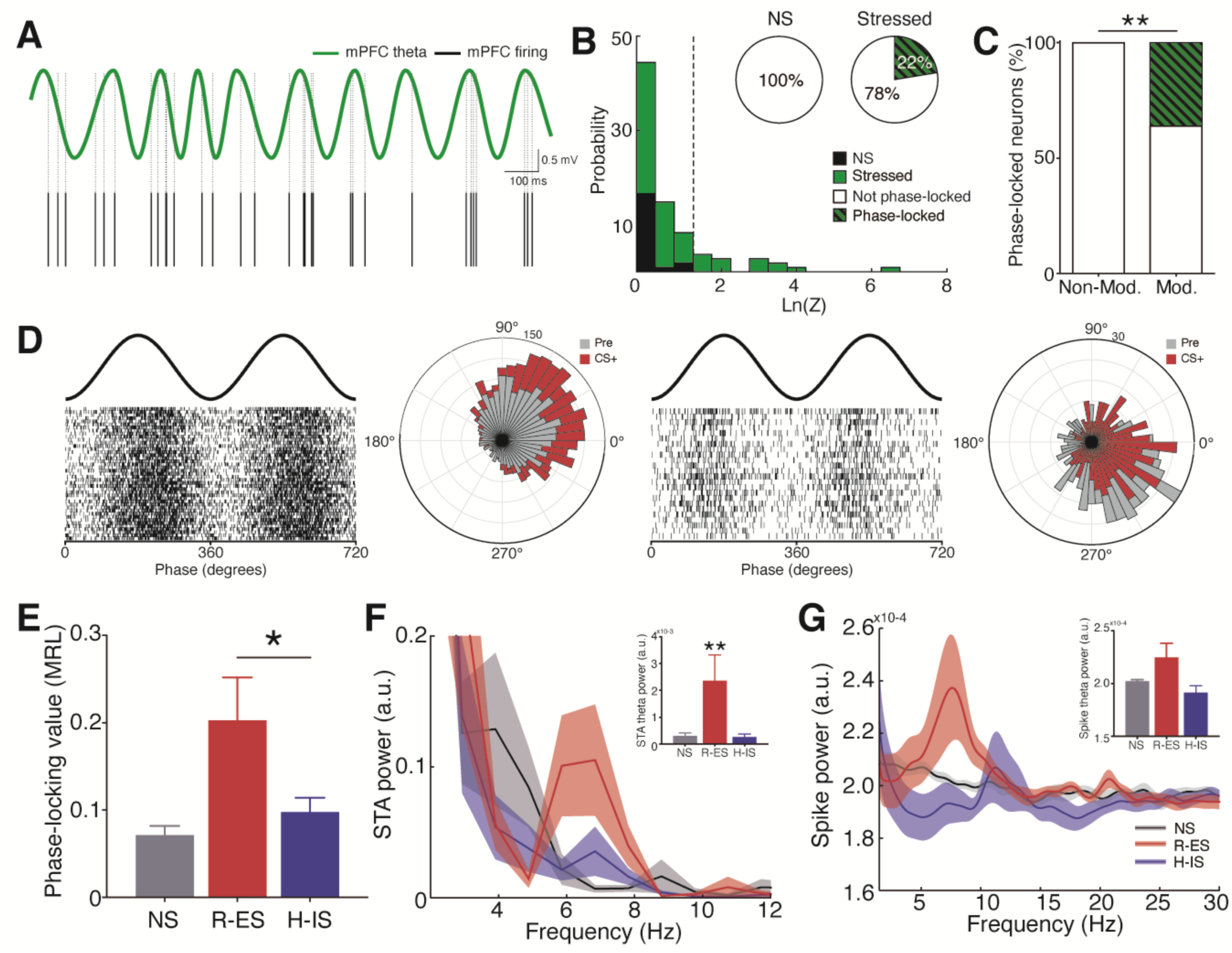
mPFC Neurons Modulated under Controllable Stress are Coupled with Theta Phase and Rhythm. (A) Representative mPFC theta-filtered signal with a neuron exhibiting phase-locked spikes. (B) Distribution of Rayleigh’s Z parameter logarithm showing greater proportion of phase-locking in animals under stress. Graded lines indicate the neurons considered significantly phase-locked (p < 0.01, Rayleigh’s test). Pie charts show greater proportion of phase-locked neurons during stress (right) than NS (left). (C) A greater proportion of phase-locked neurons from modulated units during stress compared to non-modulated units. Note that all phase-locked neurons were modulated under stress (*p < 0.05, Chi-squared test) (D) Representative phase rasters and polar histograms of two neurons from R-ES animals showing distinct modulations by CS^+^ and strong theta phase-locking to different phases. (E) R-ES modulated neurons present stronger phase-locking value (spike phases mean resultant length) than H-IS (*p < 0.05, Wilcoxon rank-sum test). (F) Stronger spike-triggered averaged LFP theta power in R-ES modulated neurons (*p < 0.05, Wilcoxon rank-sum test). (G) Spike relative power spectral density showing prominent theta power in R-ES modulated neurons, indicating theta rhythmicity of these neurons. See Figure S7. However, we found no statistical significance between averages (p = 0.18, Wilcoxon rank-sum test). MRL = mean resultant length. STA = spike-triggered average.

### CA1 and mPFC Network Activities Accurately Discriminate Individuals under Stress and Predict Resistance to Future Uncontrollable Stressors

Throughout this study, we reported a number of electrophysiological markers discriminating stressed from non-stressed animals (stressor-related variables), and resistant from helpless animals (controllability-related variables). We show that these variables are associated with distinct features of CA1-mPFC theta oscillations, which in turn coordinate mPFC neuronal firing patterns related to controllability. Our final approach was to explore patterns of activity that collectively comprise these variables and to examine how they could distinguish the effects of stress per se and predict resistance or helplessness.

The most relevant discriminators of stress and NS were: (1) elevation of CA1 theta peak frequency after US (% from CS) (AUC = 0.97; Figure S4); (2) reduction of CA1 beta ERPP (maximum at 24.0 Hz; AUC = 0.99; Figure 1, S3); (3) CA1-mPFC theta synchronization (AUC = 0.92; Figure 4) (see Figure 8A1-3i). In turn, the most relevant predictors of learned resistance and helplessness were: (4) CA1-mPFC theta synchrony during CS^+^ (AUC = 0.87; Figure 4); (5) mPFC prestimulus theta-gamma MI (AUC = 0.79; Figure 5); (6) increase in CA1 theta ERPP (maximum at 7.7 Hz; AUC = 0.92; Figure 2, S3) (7) mPFC ERPP spectrum (2.5-50 Hz) PC1 score (AUC = 0.95; Figure 3) (see Figure 8A4-7). We reasoned that we would be able to assemble the variables separately related to either stressor or controllability by computing factor analysis for two common factors (Figure 8A-B). In fact, factor 1 loadings showed greater weights at controllability-related variables (> 0.5), and lower weights at stressor-related variables (*controllability factor*) (Figure 8B). In turn, factor 2 loadings presented greater weights at stressor-related variables (> 0.5), intermediate weights at variables that were greater in both groups of stressed animals but higher in resistant ones (> 0.2), and lowest weights at variables with bi-directional modulation by the degree of control (*stressor factor)* (Figure 8B). Hierarchical clustering confirmed such distinction by showing that stressor-related and controllability-related variables are indeed deeply dissociable (Figure 8C). Thus, we obtained two scores representing the collective patterns of electrophysiological features related specifically to either controllability or stress per se.

**Figure 8.**
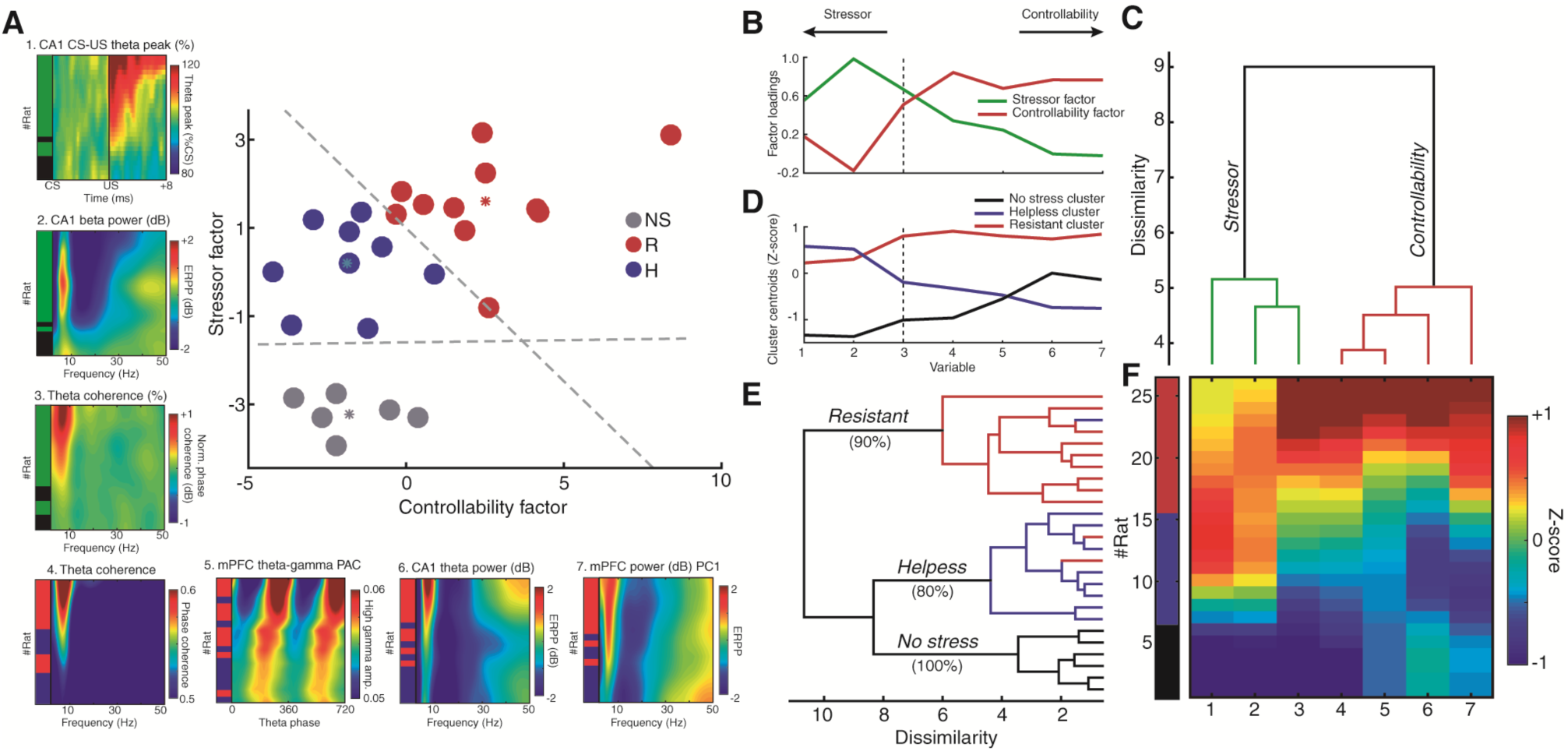
CA1 and mPFC Network Activities Discriminate Individuals under Stress and Predict Resistance. (A) A linear model accurately discriminated individuals under stress and predicted resistance. (A1-7) electrophysiological variables used in the model: (1) change in US theta peak frequency change (% from CS); (2) decrease in CA1 beta (24 Hz) power; (3) change in theta mean phase coherence (% from pre-CS); (4) Theta mean phase coherence during CS; (5) mPFC theta-high gamma phase-amplitude coupling (MI); (6) CA1 theta (7.7 Hz) power perturbation; (7) mPFC spectral power perturbation PC1 score. The left bars indicate stressed (green) vs. NS (black) individuals discriminated for variables 1-3, and R (blue) vs. H (red) for variables 4-7. *Stressor factor* and *Controllability factor* scores assemble the effects that distinguish stressed vs. NS and R vs. H respectively (see Methods and Results). Dashed lines are the functions of the regularized linear discriminant classifier model with 0.95 gamma amount of regularization. Note that although H are clearly distinct from NS by *Stressor* scores, they are identical in *Controllability* scores, indicating that these individuals showed the effects of stressor per se but lacked the controllability pattern of activity that was distinctive of R animals. (B) Stressor and controllability are two dissociable factors influencing CA1-mPFC network activity. Factor analysis for two factors returned factor loadings that weighted specifically on controllability-related variables (factor 1: *Controllability factor*) or stressor-related variables (factor 2: *Stressor factor*). Note that *Stressor factor* loadings were greater at stressor variables that did not distinguish between R and H (1-3) and were intermediate at variables that distinguished stressed from NS, and R from H (4-5). *Controllability factor* loadings were particularly greater in controllability-related variables (4-7). (C) Hierarchical clustering revealed that stressor-related (1-3) and controllability-related (4-7) variables were indeed deeply dissociable. (D to E) Hierarchical clustering revealed three distinctive clusters with specific (D) patterns of activity (cluster centroids) and that (E) predominantly comprised individuals from each behavioral category. The *Helpless* and *No stress* clusters were clustered together in relation to the *Resistant* cluster, which indicates a more distinguishable activity in R. Note that the R pattern (D, red) has both *Stressor* and *Controllability* factors, while the H pattern (D, blue) is mostly characterized by the Stressor factor, and the NS pattern (D, black) is the opposite of the Stressor factor. (F) Standardized values (smoothed Z-score) of each electrophysiological variable used in the model. Subjects were organized in NS, H and R, and were ordered by the sum of all effects.

Then we fitted a regularized linear discriminant classifier model based solely on these two predictors: stressor and controllability scores. Remarkably, our model reached 100% classification accuracy for all NS, R and H individuals simultaneously (100%, cross-validation: 92%, N = 26; Figure 8A). Additionally, the model was also able to predict the behavioral outcomes of R-IS and H-ES animals (100%, N = 6), when fitted against NS, R-ES, and H-IS data (N = 20). Furthermore, we noticed that the arrangement of individuals revealed a striking characteristic. Although *stressor scores* were greater in both R and H animals compared to NS (F(2,23) = 52.49, p < 0.0001; R vs. NS t(23) = 10.21, p < 0.0001; H vs. NS t(23) = 6.93, p < 0.0001; R vs. H t(23) = 3.37, p = 0.002), *controllability scores* were only significantly greater in R individuals (F(2,23) = 15.51, p < 0.0001; R vs. NS t(23) = 4.31, p = 0.0003; R vs. H t(23) = 4.95, p < 0.0001), while H and NS were virtually identical (H vs. NS t(23) = 0.06, p = 0.94). These findings demonstrate that resistance is associated with a collective pattern of activity that is weaker or lacking in H animals during stress.

We also examined the presence of collective patterns of electrophysiological data across individuals through hierarchical clustering. This examination revealed three clearly distinct clusters fitting almost entirely the three behavioral categories of our study: R, H and NS (X^2^(4, N = 26) = 38.86, p < 0.0001; cluster 1: 100% NS, N = 6/6; cluster 2: 80% H, N = 8/10; cluster 3: 90% R, N = 9/10). In accordance with our model, the distinguishable patterns of activity (represented by cluster centroids, Figure 8D) showed that the *resistant cluster* pattern was constituted by the summation of stressor and controllability effects, while the *helpless cluster* pattern exhibited a clear resemblance to the *stressor factor* (Figure 8D, see also Figure 8B). This finding further illustrates that the neural activity underlying helplessness is essentially characterized by the effects of stress per se. Moreover, the predominantly *helpless cluster* and the *no stress cluster* were clustered together in reference to the *resistant cluster*, which demonstrates that this latter group had the most distinctive activity (Figure 8E). In fact, we identified the most distinctive collective pattern across subjects through PCA (48% explained variance, N = 26), and we observed that the PC1 scores showed the highest univariate prediction of R individuals (AUC = 0.98, n = 11/26), and the greatest correlation with escape performance reported in this study (r(18) = 0.77, p < 0.0001; not shown).

In summary, we found a clear dissociation between the effects of controllability and stress per se that defined the unique patterns of activity of non-stressed, resistant, and helpless individuals. Altogether, our findings converge to suggest that learned stressor resistance is associated with a distinctive collective pattern of enhanced CA1-mPFC network theta activity during stress that is predominantly lacking in helpless individuals.

## Discussion

This is the first electrophysiological characterization of animals under controllable or uncontrollable stress. We evidence that stressor controllability is associated with a distinctive pattern of enhanced hippocampal-prefrontal theta activity that predicts resistance against subsequent uncontrollable shocks, and that was predominantly absent in helpless individuals. The *controllability pattern* consisted of basal theta-gamma phase-amplitude coupling and higher CA1-mPFC theta power and synchrony during CS^+^. The neuronal activity related to control was represented by a stronger phase-locking to theta rhythm and a rapid return to baseline firing after shocks. In contrast, helplessness was marked by lasting increases in mPFC firing after US, in addition to clearer CS-locked theta phase resetting. In turn, the s*tressor pattern* was characterized by a broad reduction of spectral power, CA1-mPFC theta synchronization during CS^+^, and elevated theta peak frequencies after footshocks (see Table 1).

**Table 1.** Neural correlates of stressor, controllability and uncontrollability. * = variables used in the linear model, CA1 = hippocampal Cornu Ammonis, mPFC = medial prefrontal cortex, CS = conditioned stimulus (light cue), US = unconditioned stimulus (footshock).

| Variable | STRESSOR | CONTROLLABILITY | UNCONTROLLABILITY |
| --- | --- | --- | --- |
| CA1 theta power* |  | ↑ | ↓ |
| CA1 wide range power* | ↓ |  |  |
| mPFC theta power |  | ↑ | ↓ |
| mPFC spectral perturbation pattern* |  | ↑ | ↓ |
| CA1 theta peak frequency* | US>CS |  |  |
| CA1-mPFC synchrony** | ↑ | ↑↑ |  |
| mPFC theta phase resetting |  |  | ↑ |
| mPFC theta-gamma coupling* |  | ↑ |  |
| mPFC firing rate modulation | ↑ | CS>US | US>CS |
| mPFC neuronal phase locking | ↑ | ↑↑ |  |

Experimental models of stress vulnerability usually count on natural individual predispositions to either susceptible or resilient phenotypes (Feder et al., 2009; e.g. Bagot et al., 2016; Christensen et al., 2011; Hultman et al., 2018; Krishnan et al., 2007). Although fruitful, this approach seldom considers predisposing factors that prone individuals to distinct consequences of stress, making it challenging to link biological findings to specific variables. In contrast, the triadic design of learned helplessness discriminates the effects of the degree of control over stress, a well-established modulator of behavior (Eisenstein and Carlson, 1997; Maier and Watkins, 1998; Weiss, 1968), neurochemistry (Amat et al., 2005; Bland et al., 2003; Weiss et al., 1981), circuit activation (Baratta et al., 2009, 2019), and neural plasticity (Christianson et al., 2014; Shors et al., 1989, 2007). We contribute to such a framework by reporting the effects of stressor controllability on network dynamics.

Our data link R-ES and H-IS to well defined neural signatures, which can now be interpreted as correlates of the degrees of control over stress. Also interestingly, the neurophysiological profiles of the unforeseen minor subsamples R-IS and H-ES were consistent with their behavioral outcomes rather than their programmed stress exposure. We can attribute the incidence of R-IS to individual predispositions or accidental contingencies during the experiment (e.g., Skinner, 1948). In turn, the absence of the controllability signature in H-ES indicates that this activity is more linked to the “immunization” effect of behavioral control than escape responding. In fact, mPFC inhibition has been shown to spare escaping behavior, but to abolish the protective effects of controllability against subsequent induction of helplessness (Amat et al., 2005, 2006).

### Theta Functions and Stressor Controllability

Mounting evidence indicates that theta oscillations have a role in the processing of aversive information (Bocchio et al., 2017; Çalışkan and Stork, 2019; Gray and McNaughton, 2000; Likhtik and Gordon, 2014). Many reports describe increased theta power and synchrony during different stressors, including exposure to distant predators, aggressive conspecifics, anxiogenic environments, and conditioned fear (Adhikari et al., 2010; Hultman et al., 2018; Lesting et al., 2011; Mikulovic et al., 2018; Sainsbury et al., 1987; Seidenbecher et al., 2003). These observations suggest that theta oscillations may signal aversiveness and represent a correlate of fear and anxiety. In our study, we observed increases in theta power in resistant, but not helpless animals. This finding is intriguing, given that helplessness is linked to exaggerated passive fear responses (Baratta et al., 2007). In this context, Courtin et al., (2014) reported suppression of mPFC theta power during freezing, whereas Karalis et al., (2016) showed that abolishing theta activity does not alter freezing behavior. Although we observed CS^+^-evoked theta synchronization in both resistant and helpless animals – which supports its role in signaling aversiveness – the strength of this synchrony was related to escape performance, which favors the role of theta activity in active coping. Adhikari *et al.* (2010) also reported that increased CA1-mPFC theta synchrony in anxiogenic places predicts active avoidance toward safer zones. Thus, high theta states during immobile fear behavior could be interpreted as the encoding of active responses, which is supported by the role of theta oscillations in sensorimotor integration and action selection (Bender et al., 2015; Oddie and Bland, 1998).

In this regard, we show that the striking differences in network dynamics related to stressor controllability only occurred during the CS^+^ period. On the other hand, we found consistent US-induced increases of theta frequency regardless of controllability. This finding is consistent with reports showing that theta frequency increases with movement but persist for some time after the animal ceases it (McFarland et al., 1975), e.g., after high shock-avoidance jumps (Lenck-Santini et al., 2008; Vanderwolf, 1969; Whishaw and Vanderwolf, 1973) or pain-evoked behaviors (Khanna, 1997; Tai et al., 2006). Noteworthy, we restricted our analyses to locomotion-free CS^+^ epochs. Thus, the neural signatures we found during the anticipation of shocks may indeed represent correlates of distinct “expectations” of either controllability or uncontrollability. Finally, another remarkable finding reported here was the correlation of mPFC theta power-synchrony during stress with later escape performance. In fact, extensive research has demonstrated that theta activity correlates with numerous forms of cognitive performance mostly related to memory encoding and retrieval, and executive functioning (Cavanagh and Frank, 2014; Fuentemilla et al., 2014; Hasselmo and Stern, 2014; Korotkova et al., 2018). Taken together, our results suggest that theta activity supports cognitive mechanisms of stress resistance.

The intermediate CA1 sends projections to the mPFC and has been shown to participate in both cognitive and emotional functions (Burton et al., 2009; Fanselow and Dong, 2010). Theta oscillations are generated in the septal-hippocampal circuitry and synchronize with downstream targets under cognitive demand (Buzsáki, 1996, 2002; Gordon, 2011; Harris and Gordon, 2015; Vertes and Kocsis, 1997). Indeed, we observed that CA1 theta field entrained the mPFC with a constant lag of 49 ms, consistently with previous reports (Siapas et al., 2005). However, we present evidences that local mPFC theta activity is important for stressor controllability. We observed a stronger correlation of mPFC theta power with escape performance, and a unique association between stress resistance/controllability and mPFC theta coupling to both fast oscillations and neuronal firing – which are known to be locally restricted (Buzsáki et al., 2012). Moreover, we report that differential firing to CS^+^ and US possibly encode controllability-related information, and we evidence that these firing patterns are coordinated by theta rhythm. Prelimbic neurons have also been shown to signal aversiveness (Adhikari et al., 2011; Burgos-Robles et al., 2009; Courtin et al., 2014; Diehl et al., 2018; Sotres-Bayon and Quirk, 2010). Thus, we can interpret the immediate return to baseline firing after controllable US to represent the realization that aversiveness is no longer present. In contrast, the enduring responses after uncontrollable shocks in helpless animals would represent an impairment in such realization. It is worth mentioning that animals tend to generalize the expectations of either controllability or uncontrollability (Maier and Seligman, 1976), which makes it challenging to attribute single-neuron responses to distinct degrees of control. Nevertheless, our findings are sufficient to indicate that theta oscillations coordinate local mPFC activity associated with the encoding of stressor controllability.

### Theta Impairment and Helplessness

Our study suggests that helplessness might be associated with impaired theta engagement. This is consistent with reports that uncontrollable stress induces brain-wide serotonergic activation, which is the main modulatory system that suppresses theta (Maier and Watkins, 1998; Puig and Gener, 2015; Vertes and Kocsis, 1997). Serotonin-induced effects related to helplessness arise mostly from the dorsal raphe nucleus (Maier and Watkins, 2005), which receives regulatory projections from the mPFC (Pollak Dorocic et al., 2014). This descending pathway has been consistently shown to mediate the protective effects of behavioral control (Maier, 2015; Warden et al., 2012). Furthermore, although there are no known projections from the hippocampus to the dorsal raphe nucleus, its neurons can be phase-locked to the theta field (Kocsis and Vertes, 1992; Pollak Dorocic et al., 2014). Thus, we speculate that theta influence on the dorsal raphe nucleus may be mediated via mPFC, and that theta synchrony between these two regions may facilitate brain-wide stress-protective effects. Future studies should address these questions.

Maier and Seligman initially proposed that *uncontrollability* would be the key variable that, once learned, would change behavior (Maier and Seligman, 1976; Seligman and Maier, 1967). Decades later, when mPFC inhibition during controllable stress was shown to result in helplessness (Amat et al., 2005), the authors revisited the theory to suggest that actually *controllability* is the determining factor to be learned. In this sense, helplessness would develop as a default response to severe stress if controllability is absent (Maier and Seligman, 2016). We found that resistant and helpless individuals share electrophysiological markers of stress, but only resistant animals present the neural signature of controllability, in line with Maier and Seligman (2016). Considering that theta oscillations regulate synaptic plasticity and learning (Bocchio et al., 2017; Buzsáki, 2002; Hasselmo and Stern, 2014), theta activity impairments in helplessness could also mean that this syndrome stems from learning deficits rather than a learned response.

Taking into account the translational validity of helplessness, our findings add to growing evidence that successful treatments for depression are associated with enhanced frontal theta in humans. Increased prefrontal theta predicts response to placebo (Leuchter et al., 2002) and deep brain stimulation (Broadway et al., 2012; Widge et al., 2019). Ketamine – which induces theta-gamma coupling (Ahnaou et al., 2017; Caixeta et al., 2013) – and prefrontal theta-burst magnetic stimulation (Chung et al., 2015; Li et al., 2014) show rapid effects against depression (Kraus et al., 2019; Krystal et al., 2019). Together with this literature, our findings highlight that changes in prefrontal theta may help guide therapeutic developments and approaches.

### Conclusions

Theta oscillations have been discussed to signal states of fear, anxiety, and stress vulnerability. By adding the controllability dimension to this scenario, we show that CA1-mPFC theta activity may actually play a role in stress resistance. In light of our findings, we propose that the functions of hippocampal-prefrontal theta in stress, aversion, action selection, top-down regulation, learning, and cognitive control are integrated into a multidimensional continuum that underlies active coping against stressors.

## Acknowledgments

We thank Luis Fernando Luca, Antonio Renato Meirelles e Silva, Renata Caldo Scandiuzzi, and Daniela Ribeiro for technical support. We thank Cláudia Maria Padovan, Maria Helena Hunziker, Frederico Graeff, and Nuno Sousa for discussions. This research was funded by the National Council for Scientific and Technological Development (D.B.M.: 134102/2013-4; J.P.L.: 423977/2016-4), the Coordination for the Improvement of Higher Education Personnel (D.B.M.: 88882.328283/2019-01), and the São Paulo Research Foundation (R.N.R.: 2018/02303-4; L.S.B.J.: 2012/06123-4; M.T.R.: 2011/04467-5; J.P.L.: 2016/17882-4).

## Author Contributions

DBM and RNR conceived the study. All authors developed the methodology. DBM and MTR conducted the experiments. DBM and RNR analyzed LFP data. DBM, RNR and LSBJ analyzed SUA data. RNR, LSBJ, and JPL supervised the project. All authors wrote and revised the manuscript.

## Methods

### Subjects

Adult male Wistar rats (8-10 weeks old, Ribeirão Preto, Brazil) were housed singly in bedded cages in a controlled-temperature room (22 ± 2 °C) on a 12 h light/dark cycle (lights on at 7 a.m.) with *ad libitum* access to food and water. The procedures followed the National Council for the Control of Animal Experimentation guidelines and were approved by the local Committee on Ethics in the Use of Animals (Ribeirão Preto Medical School, University of São Paulo, protocol 156/2014).

### Electrode Implantation Surgery

Animals were anesthetized with ketamine and xylazine (respectively: 50 mg/Kg and 25 mg/Kg intraperitoneal, followed by 70 mg/Kg and 35 mg/Kg intramuscular). Body temperature was kept constant during the entire procedure by a heating pad (37 ± 1 °C).

Chronic head caps consisted of a bilateral pair of eight-channel connectors (Omnetics): seven channels for each mPFC, one channel for each CA1. mPFC electrodes consisted of microwire bundles (teflon-coated tungsten, 50 µm) into both prelimbic areas (ventral: 3.2 mm, posterior: 3.0 mm, lateral: ±1 mm; bregma-referenced coordinates) (Bueno-Junior et al., 2018). In turn, intermediate CA1 areas were each implanted with a monopolar electrode (Ruggiero et al., 2018) (Figure S1). In addition to the electrodes, microscrews were fastened into the skull, including a ground reference in the interparietal bone area. Electrodes and screws were then cemented together with acrylic resin. Analgesic, antipyretic, and antibiotic drugs were injected after surgery. Animals were allowed to recover for 8-9 days before stress exposure.

### Apparatus

We customized a shuttle box system for simultaneous electrophysiological recording, shock delivery, and behavioral monitoring. The system included a relay switch between the recording cable and the preamplifier (see Extracellular Recordings). Based on pilot tests, automatically turning off this switch during footshocks (at millisecond precision) was proven necessary to avoid grounding through the recording cable, thus assuring consistency of shock intensity. The behavioral apparatus was located inside a soundproof box, and consisted of a chamber (54 cm length x 50 cm height x 33 width) divided in two compartments by a removable wall (1 cm length x 1.5 cm height). Footshocks were delivered through stainless steel bars on the floor (see Behavioral Protocol).

### Behavioral Protocol

All behavioral procedures were performed during the light phase (8 a.m. to 6 p.m.) in a controlled-temperature (22 ± 2 °C) dark room (0 lux). On day 1, rats were divided into three groups according to the triadic design of behavioral immunization (based on Amat et al., 2006). Animals underwent escapable shocks (ES), yoked inescapable shocks (IS), or no shocks (NS) (Figure 1A). ES animals were submitted to 100 trials consisting of a conditioned stimulus (CS^+^, LED light, 200 lux, 10 s fixed duration), immediately followed by the unconditioned stimulus (US, footshock, 0.6 mA, 10 s maximum duration, unless terminated by escape behavior). Inter-trial intervals were in the range of 40 ± 20 s, randomly. ES animals were allowed to escape by jumping the short wall between compartments. Yoked IS animals were exposed to footshocks of equivalent intensity and durations, but could not control them by wall jumping. NS animals were exposed only to the CS^-^. On day 2, both ES and IS animals were exposed to 40 trials of uncontrollable IS of 10 s fixed duration. On day 3 (the test session), resistant or helpless behaviors were determined by evaluating escape performance across 30 trials of ES (at fixed ratio 1) in the same shuttle box apparatus (adapted from Joca et al., 2003), but without the wall (Figure 1A). Behavioral responses were recorded by an automatized software, and all sessions were video monitored.

To classify resistant versus helpless individuals, we used k-means clustering (three clusters) with two behavioral dimensions from the test session: mean latency to escape and total number of failures (Figure 1D) (Chourbaji et al., 2005; Vollmayr and Henn, 2001; Wang et al., 2014). All NS animals were included in the cluster with best escape performance. ES or IS animals that were clustered together with NS were labeled as resistant (R), while all others in the poor-performance clusters were labeled as helpless (H) (Figure 1D). Other statistical criteria based on NS behavior, such as the maximum, mean plus two standard deviations, and Tukey’s fences outlier range returned the same results, confirming the robustness of our behavioral labeling.

### Extracellular Recordings

Electrophysiological signals were recorded during behavioral sessions, interrupted only during footshocks (see Apparatus). A multichannel acquisition processor (Plexon) was used to record local field potentials (LFP) and multi-unit activity (MUA) with the following parameters. LFP: 1000x gain, 0.7-500 Hz band-pass filter, 1 kHz sampling rate. MUA: 1000x gain, 150-8000 Hz band-pass filter, 40 kHz sampling rate. Timestamps were acquired from the behavioral apparatus at 40 kHz sampling for peristimulus analyses.

### Histology

Immediately after the test session, animals were euthanized with CO_2_ asphyxiation, and electrolytic lesion currents (1 mA) were applied between pairs of wires. After decapitation, each brain was placed in a cassette for immersion in 4% paraformaldehyde overnight (PFA, −20 °C), followed by 70% ethanol, and paraffin for coronal sectioning at the microtome. Standard cresyl-violet staining was used to validate electrode positioning under the bright-field microscope. Based on histology, we excluded 1 animal from all analyses, and 1 animal from CA1 analysis. Other 5 animals were excluded due to electrical noise or excessive locomotion.

### Data Analysis

Signal processing and analysis of electrophysiological data were performed using custom scripts in MATLAB.

### Local Field Potentials

LFP were epoched in two peristimulus windows: 8 s around CS onset (4 s pre-, 4 s post-CS onset) (Figures 2, S2, S5, 3, 4, and 5), or 24 s periods encompassing CS and US (8 s pre-CS, 8 s during CS, 8 s after US) (Figure S4). Post-US epochs, more specifically, started always 2 s after US offset. Signal was band-pass filtered (1-250 Hz), and epochs with locomotion were excluded based on power spectrum saturation and video inspection. Remaining epochs were then subtracted by the averaged epoch. Frequency bands were determined as delta (1-4 Hz), theta (5-10 Hz), alpha/beta (10-30 Hz), low gamma (30-50 Hz), and high gamma (80-110 Hz). For theta band-pass filtering we used a broader band of 4-10 Hz (Buzsáki, 2002; Buzsáki and Draguhn, 2004).

Power spectral density (PSD) was calculated using Welch’s method (1 s Hamming windowing, 90% overlap, 8192 points). Relative PSD was calculated by dividing the mean PSD estimates by the sum of the averaged pre-CS PSD below 50 Hz (Figure 2B, S4). We also investigated theta power and peak frequency (Figure S5) by concatenating all locomotion-free 1 s time windows with 90% overlap and calculating relative PSD to the sum of averaged pre-CS PSD. Normalized PSD was obtained by the ratio in dB (*10*log10*), with the averaged pre-CS PSD for each frequency bin. We detected power peaks within 5-10 Hz per time sample across stimuli. Theta relative and normalized power (dB), and normalized peak frequency (% from the CS period) were compared.

Peristimulus time-frequency decomposition was calculated using complex Morlet wavelet convolution, with 3-20 linearly spaced cycles from 1 to 120 Hz. Full epoch dB single-trial correction was used, and the mean event-related power perturbation (ERPP) was calculated using dB normalization against the pre-CS period, as described in Grandchamp and Delorme (2011) (Figure 2-3, S2-3). For event-related potentials (ERP), intertrial coherence (ITC), and phase resetting analyses, data were not subtracted by trial averages. ERP was obtained by averaging LFP across trials (Figure S5D). ITC was calculated as the mean resultant length (MRL) of phase differences for each time sample across trials using complex Morlet wavelet (as described above) for frequencies below 30 Hz (Figure S5A). For comparisons, we used theta ITC in the initial 300 ms after CS onset. For theta phase resetting, we obtained the cosine of the linear interpolation between 0’s and pi’s assigned to the peaks and valleys of the theta band-pass filtered signals (adapted from Courtin et al., 2014) (Figure S5B). Then we calculated the intertrial variance of these amplitudes (Figure S5E). We compared the mean intertrial variance across 0.1 s bins, and in the initial 150 ms after CS onset.

Cross-structural LFP synchrony, an indicative of functional connectivity, was estimated through phase coherence (Lachaux et al., 1999). Spectral mean phase coherence (MPC) was estimated by the MRL of phase differences between signals through Welch’s method (same parameters of PSD analysis) (Figure 4D). Time-frequency decomposition was estimated by the multitaper method using 5 tapers with time-half bandwidth product of 3, in 1 s segments with 90% overlap. Time-frequency phase coherence perturbation was calculated as described for ERPP (Figure 4F). Taking into account the possible nonlinearity of theta oscillations, we also obtained the MRL of the differences between instant phases estimated through Hilbert transform of the theta (4-10 Hz) band-pass filtered signals (Figure 4C, E). To estimate the lag between signals, we calculated the cross-correlation of the theta band-pass filtered signals and located the time of correlation peaks. With this we obtained both the distribution of lags and the average cross-correlation coefficients (Figure 4A-B).

Cross-frequency phase-amplitude coupling indicates how the phases of slower oscillations modulate the amplitude of fast oscillations (Ruggiero et al., 2018; Tort et al., 2008, 2009). We estimated phase-amplitude coupling strength across pairs of frequencies by computing the modulation index (MI, as described by Tort et al., 2010). Comodulation maps were constructed for frequency band pairs varying: (1) from 1 to 50 Hz (0.5 Hz bins) in steps of 1 Hz for phase modulating; and (2) from 10 to 120 Hz (1 Hz bins) in steps of 5 Hz for amplitude modulated (Figure 5B). MI between theta band (4-10 Hz) and high gamma band (80-110 Hz) were then obtained for statistical comparisons. We concatenated and used all locomotion-free pre-CS and CS periods (5 s) rather than the entire epochs because we did not see evidence for CS-related MI perturbation.

### Single-Unit Activity

Spikes were sorted semi-automatically (Offline Sorter, Plexon). Individual neurons recorded by more than one channel were identified via cross-correlation (NeuroExplorer, Nex Technologies), and in such cases, only the spike train with the largest waveforms were included in the analysis. The samples of single units (NS: 21, R-ES: 33, H-IS: 35, H-ES: 7, R-IS: 14) were then analyzed in 30 s peristimulus epochs: 10 s pre-CS, 10 s during CS, and 10 s after US offset (similarly to LFP epoching) (Figure 6A).

Neurons were labeled as *modulated* or *non-modulated* based on stimulus reactivity through comparing pre-CS spike counts versus during-CS and post-US spike counts via one-tailed paired t-tests. *Modulated* neurons were then classified as *excited* (higher spike count) or *suppressed* (lower spike count) by each stimulus (CS or US) (Bueno-Junior et al., 2017, 2018; Wood et al., 2012) (Figure 6B). The *p*-value of this categorization was Bonferroni corrected based on the total number of neurons from all groups (N = 110, p = 0.0004) (Figure 6B). Firing rate modulation was estimated for each neuron by Z-score normalization against the pre-CS period in bins of 100 ms (Figure 6A). The same bin size was used for peristimulus time histograms. We used the mean Z-score of the initial 1 s after CS onset and initial 1 s after US offset for group comparison (Figure 6C-D).

We then explored more deeply the relationships between CS- and US-evoked responses across neurons. First, all binned Z scores during CS and post-US (i.e., their entire 10 s) were separately summed up, generating one CS and one US modulation value per neuron. Then, we calculated: (1) net difference as the difference between US and CS values (Figure S6A-B), (2) absolute difference as the modulus of the difference between US and CS values (Figure S6D), and (3) relative difference as the difference between the moduli of US and CS values (Figure 6E). To identify temporal patterns of neuronal modulation in behavioral categories (stressed, R and H) we used principal component analysis (Figure 6F-G). PC1 coefficients were obtained from mean Z-scores per bin of all neurons. For each behavioral category we calculated correlation coefficient moduli between Z-scored firing rate for each neuron and PC1 coefficients. This procedure is based on a previously described method (Chapin and Nicolelis, 1999; Kim et al., 2017; Narayanan and Laubach, 2009), with one adaptation: we replaced the PC1 strength (modulus of the sum of PC1 coefficients projected onto the Z-scored data) by the correlation coefficient, as we observed this coefficient to correspond better to patterns than intensity of modulation.

We also investigated the phase locking of single-unit activity to theta oscillations. Spike times were rounded to the LFP sampling rate (1 kHz), theta phases were assigned to each spike (Figure 7D), and their phase locking strength was estimated as the MRL (Figure 7E). The significance of phase-locking was calculated using Rayleigh’s Z test (Berens, 2015) with a *p*-value threshold of 0.01, and the Z parameter (Z = MRL*number of spikes^2^) logarithm was computed (Figure 7B). Only neurons with more than 50 spikes during CS in locomotion-free trials were analyzed (as described in Karalis et al., 2016). Spike-triggered average (STA) PSD was determined as the PSD of the averaged LFP across 1 s segments centered at each spike per neuron (adapted from Yang et al., 2018) (Figure 7F). Spiking rhythmicity was assessed by computing spectral estimates on binarized spike data (fire = 1, no fire = 0) (similarly to Rosenberg et al., 1989; Royer et al., 2010) (Figure 7G, S7). Relative STA-PSD, spike PSD, and time-frequency decomposition were computed within 1.5-30 Hz using 1024 points. Windowing parameters were the same as used for LFP analysis.

### Multivariate Analysis and Classification Model

To estimate the classification accuracy of behavioral categories by single electrophysiological variables, we calculated the area under the receiver operating characteristic curve (AUC). We explored collective patterns of relationships among electrophysiological data through unsupervised multivariate analysis. Each variable was Z-scored before multivariate analysis (Figure 8), except for power spectra (Figure 3, S4) or binned firing rates (Figure 6, S6). PCA was computed using singular value decomposition algorithm, and PC scores were obtained through the projection (i.e., sum of the pointwise multiplication) of PC coefficients onto the original Z-scored data. Common factor analysis was computed using maximum likelihood estimates to obtain factor loadings (Figure 8B). Factor scores were in turn obtained by projecting the factor loadings, as described for PCA (Figure 8A). Agglomerative hierarchical tree clustering was computed using inner squared Euclidean distance (*dissimilarity*) and clusters were determined by threshold values of dissimilarity (Figure 8C-E).

To assess the effectiveness of neurophysiological variables to simultaneously classify the three stressor controllability-related categories, NS, R, and H, we fitted a regularized linear discriminant classifier model and estimated the classification accuracy (Figure 8A). For that, we used the MATLAB function *fitcdiscr* with gamma hyperparameter optimization. We deliberately used only two predictors to allow 2D graphical representation and interpretation of model functions. Classification accuracy was finally determined as the probability of correct category assignments considering all animals, and then through leave-one-out cross-validation.

### Statistical Analysis

We used paired t-test for within-group comparisons, one-way analysis of variance (ANOVA) for comparisons between three or more groups, and two-way ANOVA with repeated measures for comparisons across time bins, or periods. We performed Fisher’s least significant difference (LSD) test as *post hoc* analysis after ANOVA. Normality was assessed using the Lilliefors test. When at least one group in a given comparison failed the normality test, we used non-parametric alternatives: Wilcoxon signed-rank test for peri-stimulus changes, and Kruskal-Wallis test to compare three or more groups, followed by the pairwise Wilcoxon rank-sum *post hoc* test. The chi-squared test was used to compare the proportions of observations (i.e. rats or units) between groups. We calculated Pearson’s correlation between normal distributions and Spearman’s rank correlation as a non-parametric equivalent. For correlation analysis across trials, we used the false discovery rate correction by the number of trials (N = 30). We applied Fisher’s Z transform to compare coherence and correlation estimates. Data are expressed as the mean ± standard error of the mean (SEM). The significance level was set to 0.05 unless stated otherwise.

## Supplemental Figures

**Figure S1.**
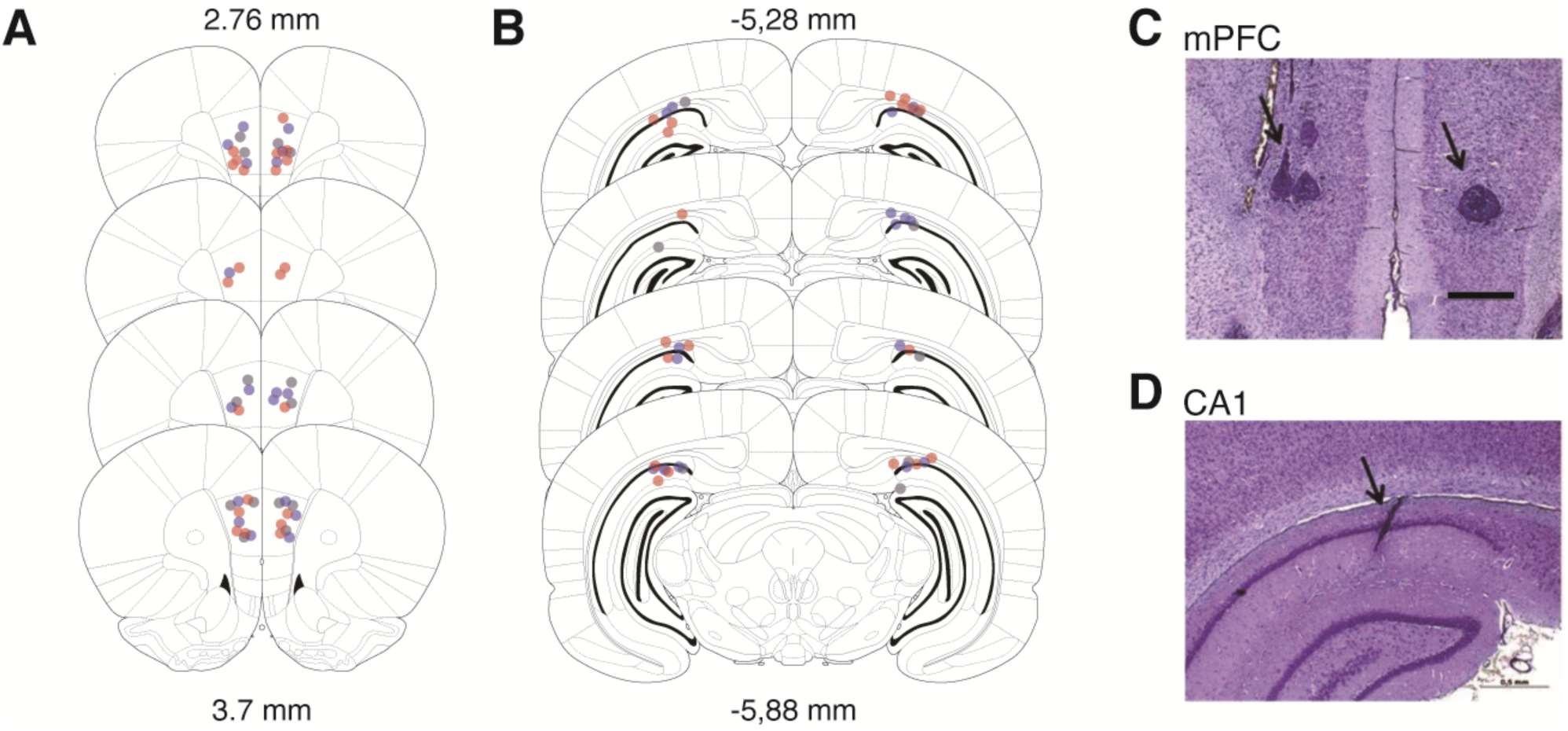
Electrode placements. (A and B) Electrode positions in the prelimbic area of the mPFC, and intermediate hippocampal CA1. Gray = NS; Blue = IS; Red = ES. (C and D) Representative electrolytic lesions.

**Figure S2.**
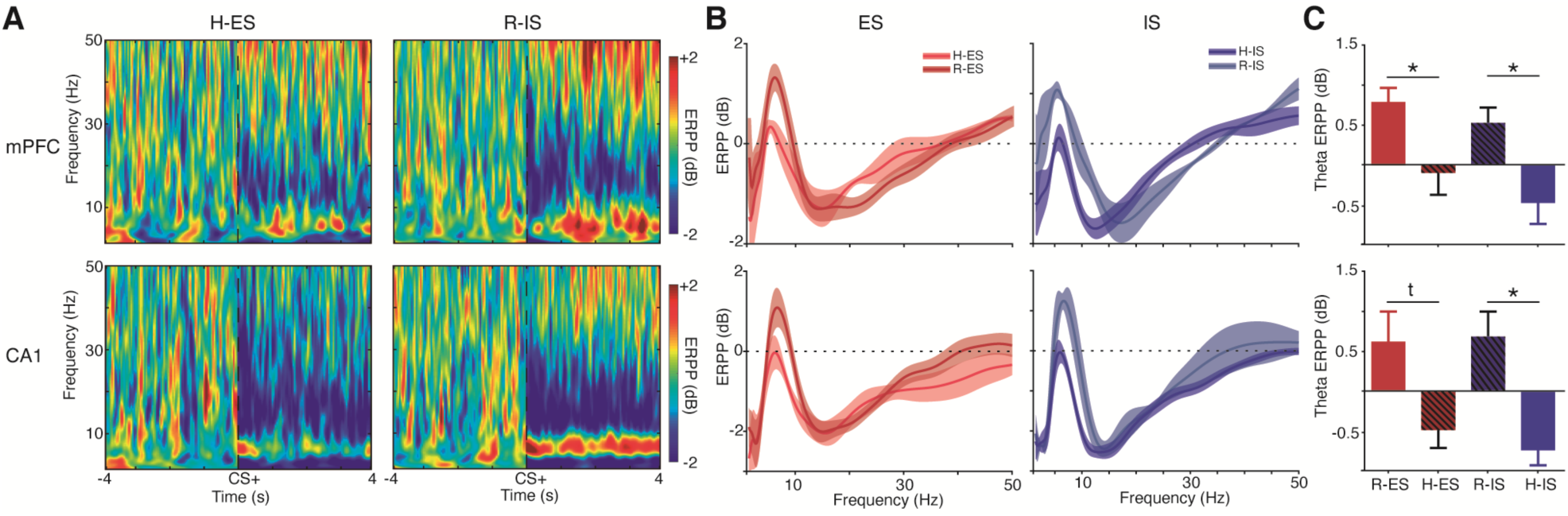
Distinctive Patterns of Power Perturbation between Resistant and Helpless Individuals during Controllable or Uncontrollable Stress. (A) Average spectrograms of mPFC (top) and CA1 (bottom) ERPP from H-ES and R-IS animals show the same profile found between R-ES and H-IS, indicating stronger association of neural activity with subsequent behavior (R or H) than with stress exposure (ES or IS). (B) Characteristic patterns of power perturbation spectrum in R and H during ES (left) or IS (right) in both CA1 (bottom) and mPFC (top). Spectra also confirm a stronger distinction within the theta band. (C) Theta power perturbation was the opposite between R and H animals, with little effects of stress exposure in both mPFC (top, F(3,16) = 7.24, p = 0.002) and CA1 (bottom, F(3,16) = 4.12, p = 0.02). Note that R-ES and H-IS showed the greatest distinction in mPFC theta power. *p < 0.05, Fisher’s LSD test.

**Figure S3.**
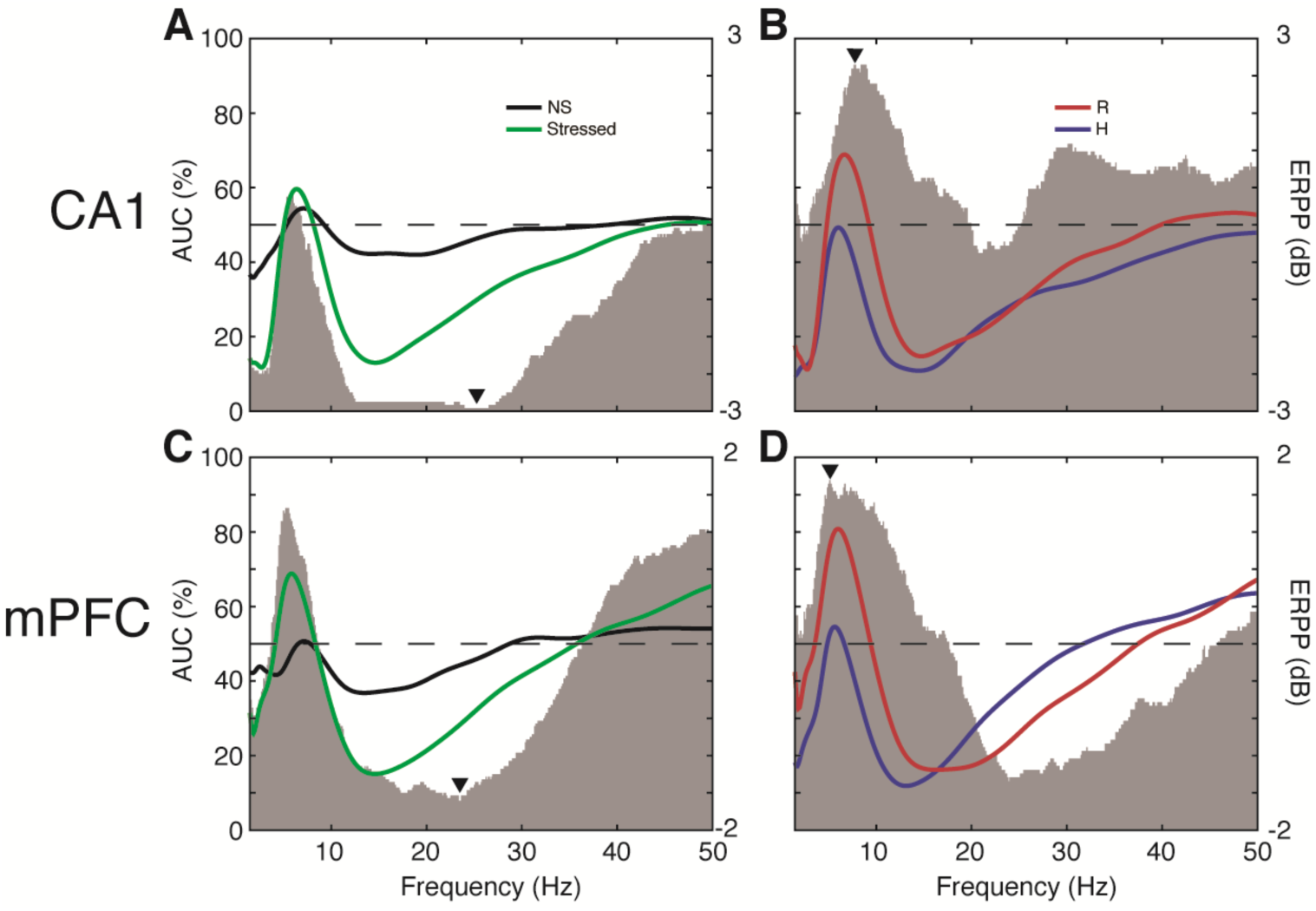
Power Perturbation Classification Performance of Stressed versus Non-Stressed, and Resistant versus Helpless Individuals. (A to D) Frequency-wise event-related power perturbation binary classification performance estimated by the area under the receiver operating characteristics curve (gray patch, left y-axes). Colored lines are the average event-related power perturbation (right y-axes). (A and C) Classification performance of animals under stress (N = 20) vs. NS (N = 6). (B and D) Classification performance of R (N = 11) vs. H (N = 9). Note that (A) CA1 achieved greater discrimination between stressed and NS groups, while (D) mPFC did so for R vs. H. Also note that (A) CA1 discriminates stressed in a wide range of slow oscillations, achieving a maximum close to the upper limits (24 Hz).

**Figure S4.**
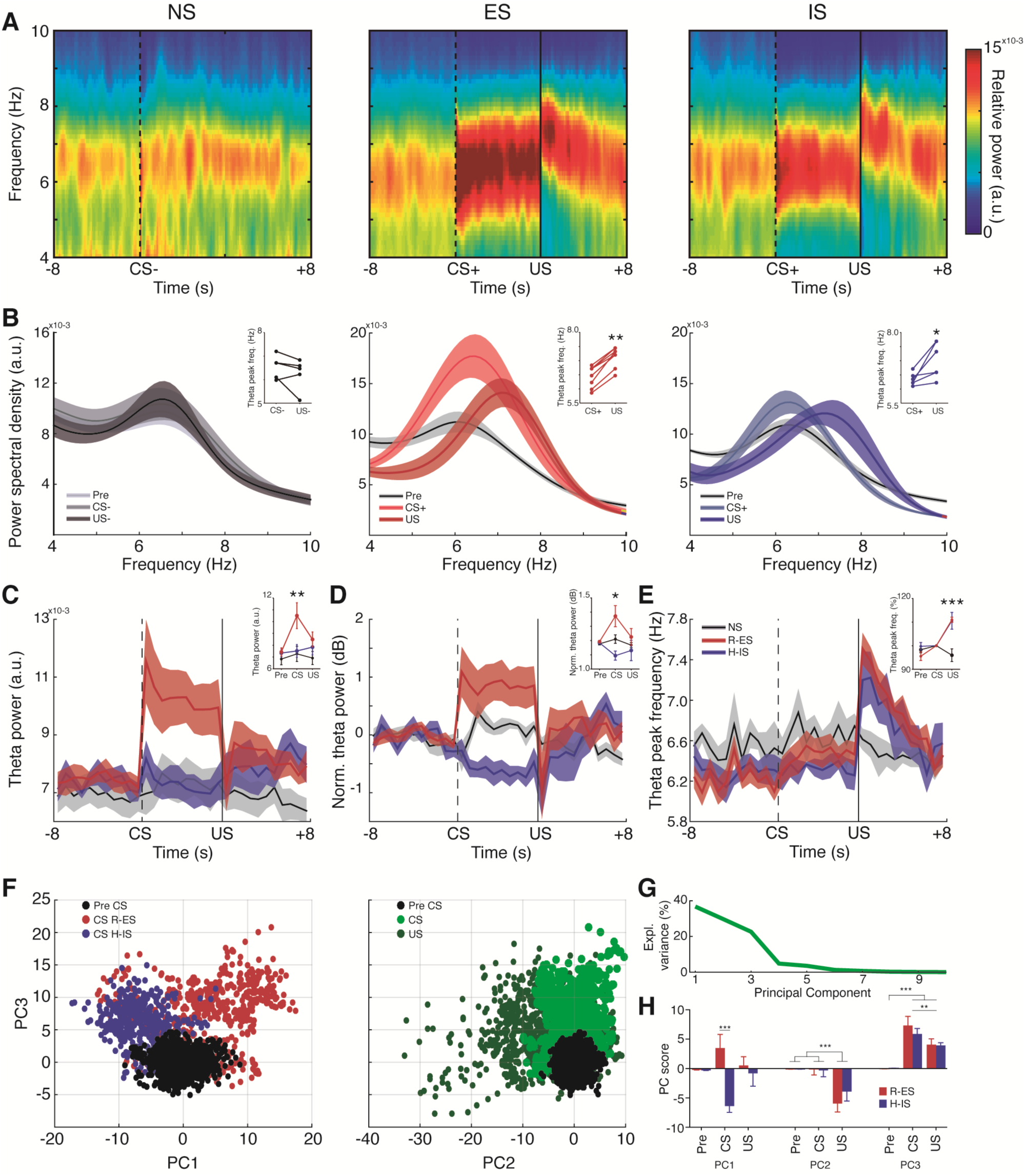
Hippocampal Theta Power and Frequency Map States of Anticipation, Reaction and Controllability over Aversive Stimuli. (A) Average power spectrograms showing that both ES and IS exhibited increased theta peak frequency after shock termination. Time samples were concatenated from locomotion-free epochs across the session. (B) Theta power spectra showing that controllability and CS^+^ modulated power, while US modulated frequency. Top right panels show increased theta frequency after US in all animals (*p < 0.05, paired t test or signed rank test). (C) R-ES presented stronger theta power only during CS^+^ (stressor controllability F(4,51) = 6.22, p = 0.003). (D) Degree of control over stress bi-directionally influenced theta power modulation during CS^+^ (stressor controllability F(4,51) = F(4,51) = 6.03, p = 0.004). (E) Theta frequency effects after US were not modulated by degree of control (period F(4,48) = 17.48, p < 0.0001). Also note that CS^+^ did not modulate theta frequency. (C to E) Insets compare the averages of 4 s periods prior to CS (pre), after CS onset (CS) and 2 s after shock termination (US). (F) PCA of normalized theta power spectra (4-10 Hz) across time samples of R-ES and H-IS individuals map states of anticipation (CS^+^), reaction (US) and controllability (R-ES vs. H-IS). (G) The three initial components of PCA presented distinctively high explained variance. (H) PC scores of each averaged period per rat distinguished particular stress-related states: PC1 discriminated R-ES and H-IS only during CS^+^ (controllability x period interaction F(2,24) = 7.29, p = 0.003) (*controllability*); PC2 discriminated US from both CS^+^ and pre-CS (period F(2,24) = 14.64, p < 0.0001) (*reaction*); PC3 mainly discriminated CS^+^ regardless of controllability (period F(2,24) = 31.02, p < 0.0001) (*anticipation*). *p < 0.05, Fisher’s LSD test.

**Figure S5.**
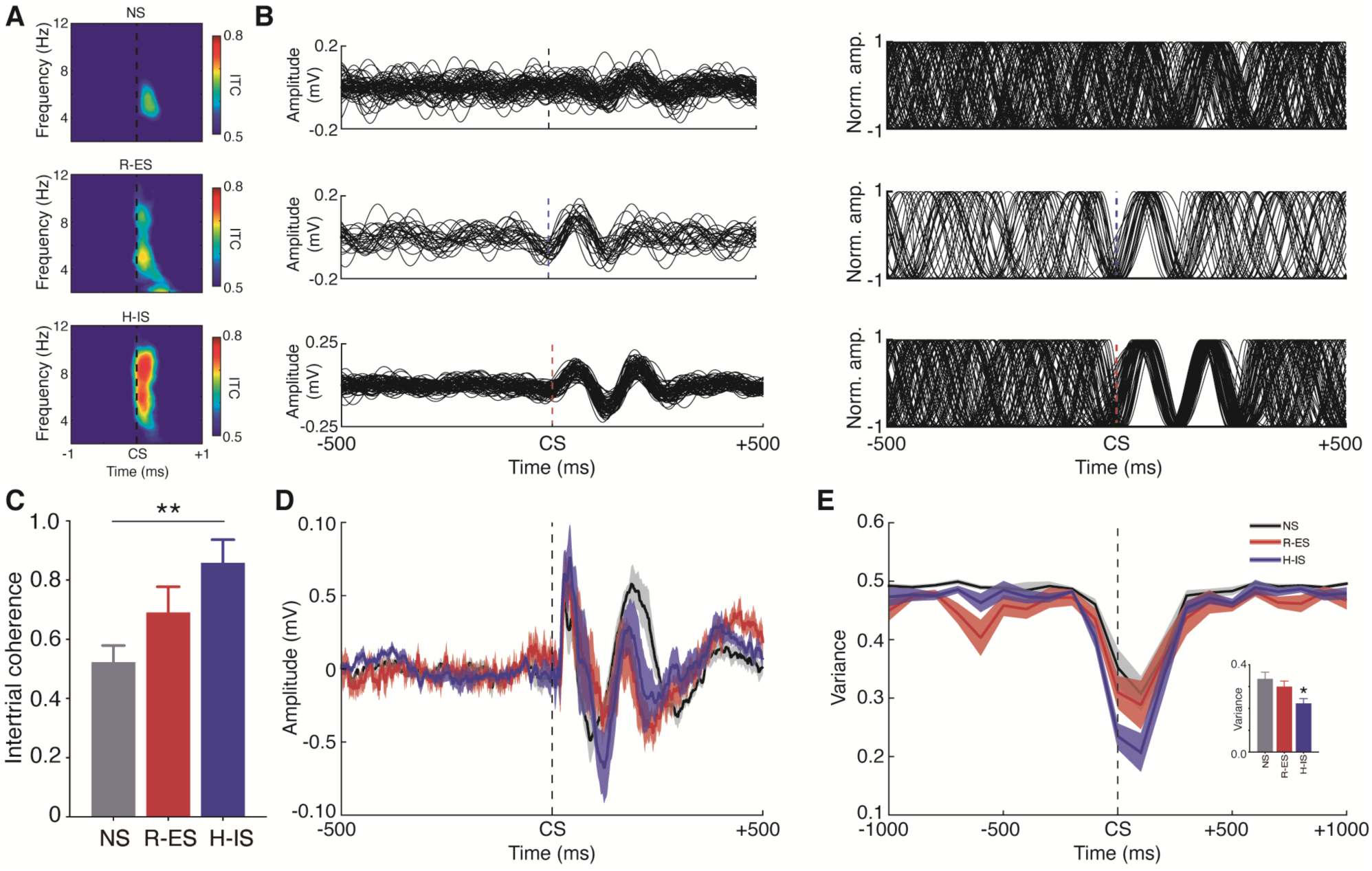
Greater CS-Evoked mPFC Theta Phase Resetting in Helplessness. (A) Average spectrograms of intertrial coherence showing a transient increase specific to the theta band in all groups. (B) Theta filtered signals (left) and standardized amplitudes (right, see Methods) across trials of representative animals from NS (top), R-ES (medium) and H-IS (bottom) showing stronger phase resetting to CS^+^ onset in the H-IS subject. (C) Greater intertrial theta coherence in H-IS than NS in the initial 300 ms (F(2,18) = 4.42, p = 0.02, *p < 0.05, Fisher’s LSD test). (D) Average event-related potentials were consistent with theta cycles. (E) Variance of the standardized theta amplitudes (see Methods) showing a CS^+^-triggered decrease in variance in all groups, especially in H-IS. Bottom right panel compares mean variances in the initial 150 ms (i.e., one theta cycle) (H(3) = 8.32, p = 0.01, *p < 0.05, Wilcoxon rank-sum test). ITC = intertrial coherence.

**Figure S6.**
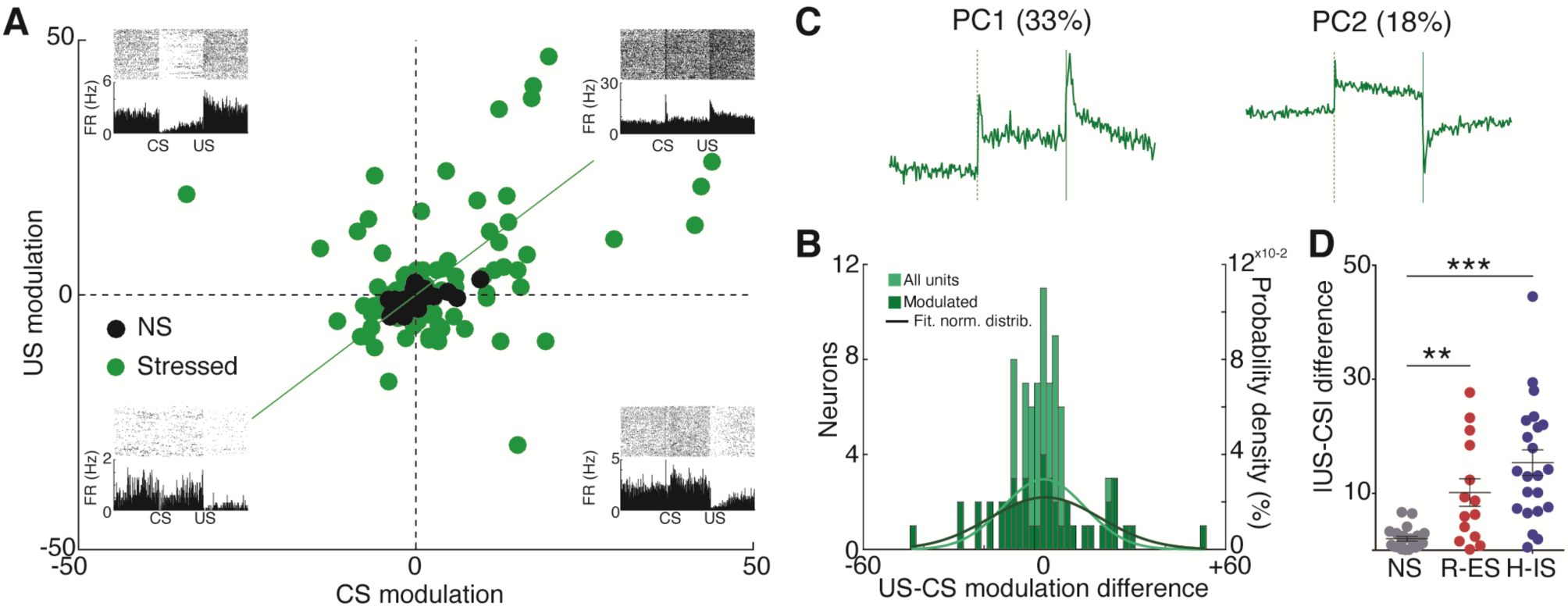
Differential Firing Rate Modulation upon CS^+^ and US during Stress. (A) Distinct CS- and US-evoked firing responses across all neurons. Each quadrant contains a representative raster plot of a neuron corresponding to that pattern. Diagonal green line represents the expected position if both modulations were equal. (B) CS- and US-modulated neurons displayed a normal distribution of US-CS differences, suggesting a role in the encoding of aversive information (Lilliefors test). The lines represent expected probability densities for normal distributions. (C) Principal components 1 and 2 from neurons of stressed animals demonstrate that the main perievent patterns are (1) stronger and (2) opposed modulation between US and CS, respectively. (D) Neurons of stressed animals exhibited higher absolute differences between Z-scored US and CS responses (F(2,54) = 14.97, p < 0.0001). The entire post-stimulus periods (10 s during CS, 10 s after US offset) were taken into account in this analysis. *p < 0.05, Fisher’s LSD test.

**Figure S7.**
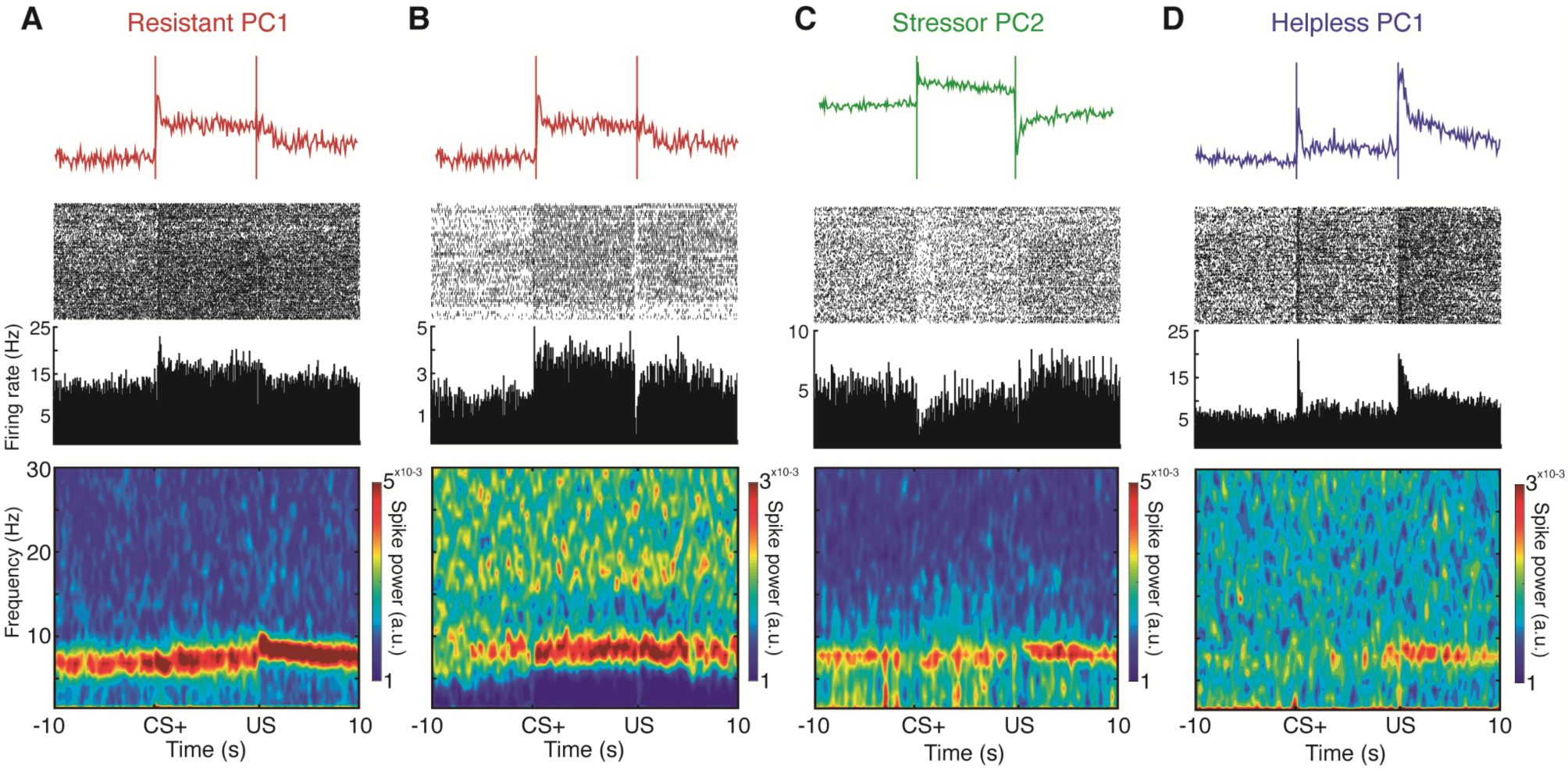
Representative Neurons exhibiting the Co-occurrence of Stressor Controllability-Related Firing Patterns and Theta Rhythm. (A to D) Representative neurons displaying perievent firing rate responses (medium) consistent with the previously described patterns revealed through PCA (top), and evident spiking theta power (bottom). These neurons indicate that neuronal representations of stressor controllability are coupled with theta rhythm. (A to C) R-ES neurons consistent with (A to B) *resistant PC1* and (C) *stressor PC2*. Note the increases in spiking theta power parallel to decreases in firing rate after US. (D) H-IS neuron consistent with *helpless PC1*. Note the weaker spiking theta power during CS^+^.

